# Phase space characterization for gene circuit design

**DOI:** 10.1101/590299

**Authors:** Macarena A. Muñoz Silva, Tamara Matute, Isaac Nuñez, Ambrosio Valdes, Carlos A. Ruiz, Gonzalo A. Vidal Peña, Fernán Federici, Timothy J. Rudge

**Affiliations:** Universidad Andres Bello, Santiago, Chile; School of Engineering, Pontificia Universidad Católica de Chile, Santiago, Chile; Department of Molecular Genetics and Microbiology, School of Biological Sciences, Pontificia Universidad Católica de Chile, Santiago, Chile; Fondo de Desarrollo de Áreas Prioritarias, Center for Genome Regulation; Millennium Institute for Integrative Biology (iBio), Santiago, Chile; Institute for Biological and Medical Engineering, Schools of Engineering, Biology and Medicine, Pontificia Universidad Católica de Chile, Santiago, Chile; Department of Chemical and Bioprocess Engineering, School of Engineering, Pontificia Universidad Católica de Chile, Santiago, Chile

**Keywords:** characterization, genetic circuit, context, transcriptional unit

## Abstract

Genetic circuit design requires characterization of the dynamics of synthetic gene expression. This is a difficult problem since gene expression varies in complex ways over time and across different contexts. Here we present a novel method for characterizing the dynamics of gene expression with a few parameters that account for changes in cellular context (host cell physiology) and compositional context (adjacent genes). The dynamics of gene circuits were characterized by a trajectory through a multi-dimensional phase space parameterized by the expression levels of each of their constituent transcriptional units (TU). These trajectories followed piecewise linear dynamics, with each dynamical regime corresponding to different growth regimes, or cellular contexts. Thus relative expression rates were changed by transitions between growth regimes, but were constant in each regime. We present a plausible two-factor mathematical model for this behavior based on resource consumption. By analyzing different combinations of TUs, we then showed that relative expression rates were significantly affected by the neighboring TU (compositional context), but maintained piecewise linear dynamics across cellular and compositional contexts. Taken together these results show that TU expression dynamics could be predicted by a reference TU up to a context dependent scaling factor. This model provides a framework for design of genetic circuits composed of TUs. A common sharable reference TU may be chosen and measured in the cellular contexts of interest. The output of each TU in the circuit may then be predicted from a simple function of the output of the reference TU in the given cellular context. This will aid in genetic circuit design by providing simple models for the dynamics of gene circuits and their constituent TUs.

---

Synthetic Biology is enabling the formalization of genetic circuit design of increasing scale and predictability^1^. The design-build-test cycle has been exemplified by the SBOL stack^2–4^ and SynBioHub^5^, which allow a standardized description for designs using a common language to represent parts and circuits. The SBOL stack and associated tools provide for creation^6^ and storage of these designs, their sequences and the experiments related to their characterization^7^ and performance, as well as models to allow simulation^8^. The designs can be linked to several repositories of parts that already exist, allowing harmonization of all the data available from these repositories in an easy way that can be shared between laboratories.

However, circuit behaviour remains unpredictable and major efforts have been focused on avoiding uncertainties that affects the design of reliable systems and their operation as specified^9^. For example Cellular Logic (Cello) is a design environment that provides automatic design of genetic circuits to perform desired operations through a set of specifications and constraints^1^. These constraints must be provided by the user and hence come from the characterization of parts and components. Thus, in order to characterize and design genetic circuits and select the parts to build them it is necessary to analyze the context in which they are introduced.

In this circuit context, we can define different levels or subcontexts. First, there is the **cellular context**, related to the physiological state of the cells which contain the genetic circuits ^10–14^. This is determined by nutrient availability and other chemical changes in the cell environment, which lead to different growth regimes, and therefore different physiological regimes. It has been shown that gene expression is strongly dependent on growth rates^15–19^, due to changes that affect the availability of resources for transcription and translation such as Polymerases, Ribosomes or dNTPs. These factors may change dynamically during the operation of a genetic circuit, for example during the growth cycle of a cell culture.

Second, we must take in count that gene circuits are carried on plasmids or inserted into the genome, where two or more Transcriptional Units (TU) can be present. These TUs consist of promoters, assembled with an RBS, a CDS and a Terminator. Inside the genetic circuit, the different TUs can interact with each other, for example when the product of one TU is capable of positively or negatively regulating another TU. Apart from these direct interactions, there is also competition between the TUs for cellular resources that allow transcription and translation^10,20–22^ and with the rest of the genes that the cell needs to express^23^ (“functional composability effects”). Further, within a circuit the spatial position inside the plasmid or genome, the orientation and the proximity of TUs can affect their expression12, 22–25 (“physical composability effects”). All of these effects constitute the **compositional context** of the genetic circuit.

Finally, the context within the TU itself is related to the parts from which it is composed, and it has been demonstrated that the flanking sequences of a part can affect its behaviour, leading to different activity of the part depending on which sequences it assembled with^27–30^. We call these effects the **sequence context**.

One of the goals of synthetic biology is to be able to characterize genetic parts to eventually predict their dynamical behaviour when used to build circuits from parts operating in different **cellular**, **compositional** and **sequence contexts**. Thus, it is necessary to develop accurate but simple models that allow characterization of parts from measurement data in a range of conditions. For example, Kelly and co-workers^31^ pointed out the importance of having standardized models for characterization and that these should be shareable between labs. They developed a method to measure promoter relative activity through analysis of the synthesis rate of GFP in different conditions. They characterized promoters relative to a reference promoter, measured separately, reducing the variance compared to the absolute promoter activity.

Later, Keren et al., 2013^32^, evaluated large libraries of promoters from *Escherichia coli* and *Saccharomyces cerevisiae* in different growth media. Using fluorescent protein fusions they showed that during maximal growth (exponential regime) promoter activities were proportional, meaning that they differed only by a scaling factor, across all conditions tested. They presented a simple resource allocation model to account for this behaviour. Effectively they demonstrated a simple linear model to characterize promoters relative to one another. This model allowed decoupling of promoter activity from **cellular context** and extraction of quantitative characteristics of promoters.

Previously we studied combinations of two promoters in the same plasmid each fused to distinct fluorescent reporters in similar TUs^33^. This allowed us to concurrently track activities of two promoters in the same cells. Similarly to Keren et al. our work showed that in exponential growth regime promoter activities were proportional, and further that their relative activities (constant of proportionality) were constant across a range of growth conditions. This model allowed us to reduce variation in promoter characteristics (relative activity) due to **cellular context** from 78% to less than a 3% of total variance.

Another approach to dealing with gene circuit context is to build systems that are designed to remove context effects. **Sequence context** has been addressed by a number of strategies: such as engineering modular insulated transcriptional elements^30^, hammerhead ribozyme and hairpin (RiboJ) structures to maintain promoter response regardless of the downstream sequence^27^, promoter insulation sequences^1,29,34^; and bicistronic devices for translation initiation (BCDs^35^). **Compositional context** due to competition for resources has been addressed using feedback control^10^ and methods to address retroactivity have been proposed by Del Vecchio et al.^36^.

These approaches require additional circuit design complexity and still may not remove all context effects^35^. Further, few of these studies analyze the time dynamics of gene expression in circuits. Thus there is a need for dynamic characterization methods that account for the various levels of context effects present in gene circuits. Here we construct simple circuits consisting of combinations of three fluorescent reporter TUs, and consider the full growth cycle of cell cultures. We track the trajectories of these circuits through the phase space of reporter expression levels. The combination of TUs and their phase space trajectories reveal the effects of **cellular** and **compositional context** on the dynamics of their expression, and suggest approaches for reliable gene circuit design that overcome them.

## Results and discussion

### Multi-fluorescent reporter plasmids for relative characterisation

In previous work we studied the relative activity of promoters of interest with respect to a common reference promoter, with each promoter in a separate TU^33^. Here we extend this work to more complex systems involving three TUs, again using one of them as a common reference. We used fluorescent reporters to construct the simplest three TU system where no TUs are explicitly coupled to any others and each TU produces a measurable output. We constructed a series of 12 plasmids from 10 TUs, each containing one of 7 promoters (Figure 1). Two of the promoters were repressible, by TetR (R0040) and LacI (R0010). Each plasmid contained three TUs; one driving RFP, another driving YFP, and a common reference driving the CFP protein. The constitutive reference TU promoter was chosen according to Kelly et al. 2009. Each RFP TU and YFP TU was designed with the same ribosome binding site with bicistronic design (BCD) translational elements^35^ (BCD2 and BCD12 respectively) and similar terminator (ECK0818 and ECK9600 respectively).

**Figure 1.**
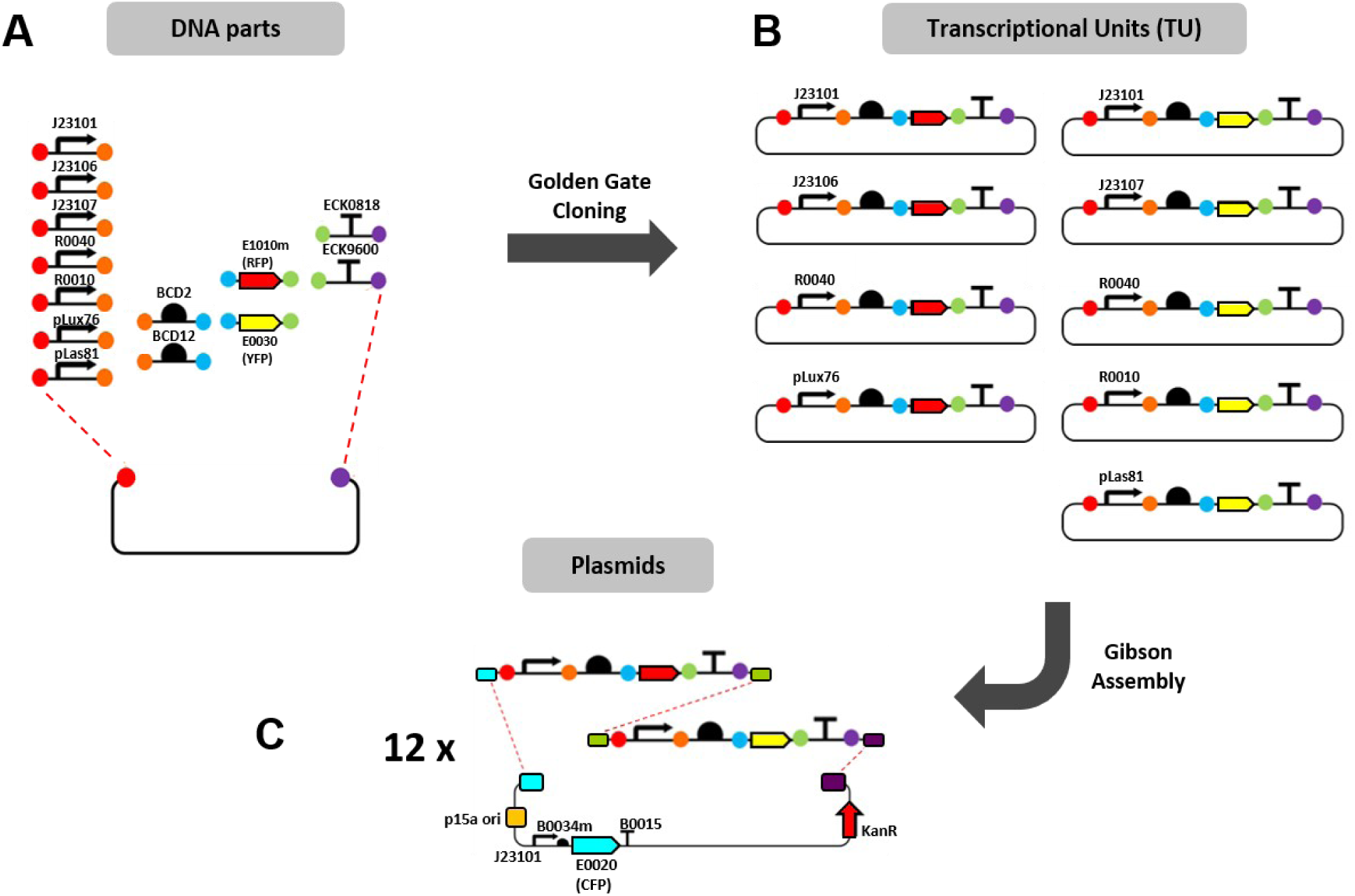
Modular assembly of plasmids. (A) Level 0 parts used for TUs assembly through Golden Gate Cloning. Each of the parts are contained in plasmids and are flanked by a restriction site and a small sequence (color circles) that allows the ordered assembly of the parts in an acceptor plasmid, obtaining a level 1 TU through Golden Gate Cloning. (B) Assembled TUs, four contained the protein E1010m (RFP) and five contained the protein E0030 (YFP). The TUs were amplified through PCR, from flanking sequences called Unique Nucleo tide Sequences (UNSes). (C) Scheme of final plasmids. The TUs were inserted in the destination plasmid containing homologous UNSes (cyan, green and purple rectangles) to the ones they contained, as well as the reference CFP TU (which has the J23101 promoter, B0034m RBS, E0020 (CFP) CDS and B0015 terminator). This allowed ordered construction of TU combinations via Gibson assembly. We obtained 12 plasmids which each contained three TUs (RFP, YFP and CFP), the p15a origin of replication (orange square) and kanamycin resistance (red arrow).

We measured the RFP, YFP, and CFP fluorescence levels, and optical density (OD) of *E. coli* containing each plasmid using a microplate reader. These assays considered the full growth process of bacteria, a dynamically changing **cellular context**, as they transition through different growth regimes over 24 hours. Repressor proteins LacI and TetR were constitutively expressed from the genome meaning that three of the TUs (R0040:RFP, R0040:YFP, R0010:YFP) were repressed during the assays.

### Phase space trajectories of plasmids reveal non-linear relationships between TUs

Each plasmid’s time dynamics form a trajectory through a 3-dimensional phase space, parameterised by RFP, YFP and CFP expression levels. To study the relative expression of RFP and YFP TUs with respect to the CFP TU over time, we projected the phase space onto the planes RFP/CFP and YFP/CFP (Figure 2). These phase space plots reveal non-linear relationships between TU expression rates, or equivalently time variation in the relative expression rates with respect to the reference CFP TU.

**Figure 2.**
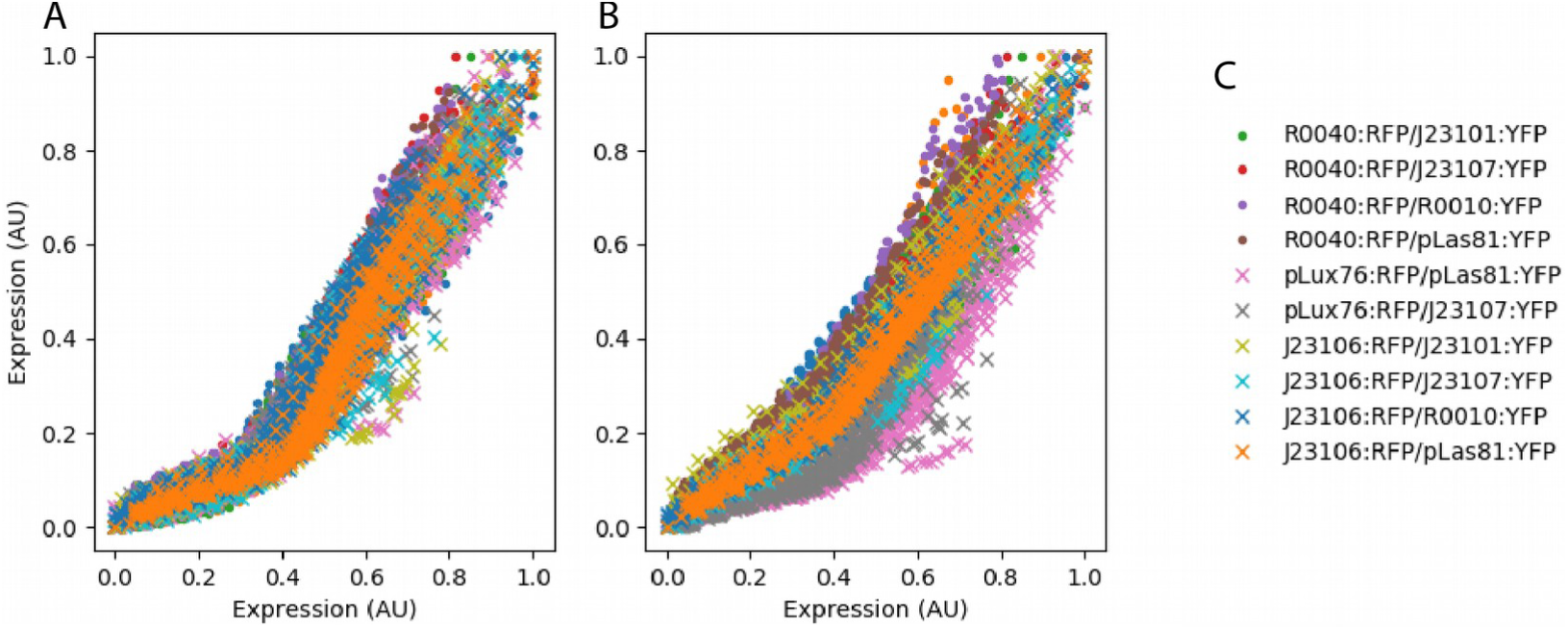
Normalized phase space plots of all plasmids showing non-linear trajectories. (A) RFP against CFP and (B) YFP against CFP, (C) legend. Fluorescence intensities are measured in AU and normalized to [0,1]. All 30 replicates for each plasmid are shown.

To understand this, consider the expression rate of a TU synthesizing a stable fluorescent protein as in this study. This can be estimated from fluorescence intensity (*I*) and optical density (*OD*) as^33^,

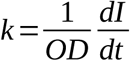

Writing the expression rates of two TUs as *k* _1_ and *k* _2_, we have the relative expression rate,

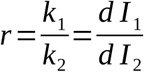

which is the slope of the phase space plot for *I* _1_ and *I* _2_, the intensities of fluorescence for each TU. We note that this approach does not require the noise-sensitive calculation of time derivatives of fluorescence nor normalisation by OD, giving a robust measure of the time dynamics of gene expression rates.

Since the slopes of the phase plots vary over the trajectories (Figure 2), we can see that relative TU expression rates are not constant, and so neither are the individual expression rates. The implication for characterisation of TUs in gene circuits is that neither their absolute nor their relative expression rate can be assumed to be constant over time varying cellular contexts.

### Phase space trajectories of plasmids follow two linear dynamical regimes

Examining the phase space trajectories of plasmids we observed that they exhibit an inflection point and can be approximated by two linear dynamics (Figure 3A). We fitted a two component piecewise linear model such that,

**Figure 3.**
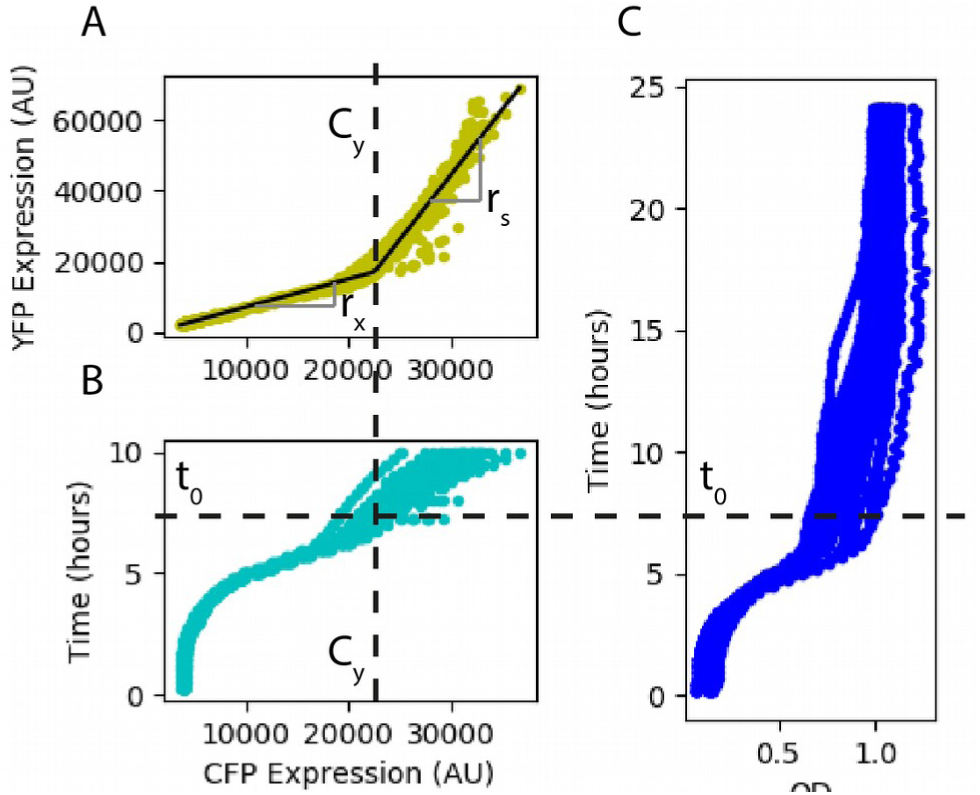
Illustration of piece-wise linear model fitting and mapping inflection point to crossing time and OD. Only YFP is shown but the same procedure was applied to RFP. (A) Fitted inflection point of YFP vs CFP, C_y_ used to find the crossing time t_0_ of the CFP time courses (B). The OD at this time (C) corresponds to the transition to stationary phase.

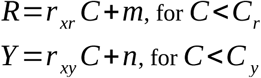

and

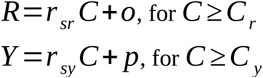

where R, Y and C are the fluorescence intensity of RFP, YFP and CFP. The constants *m, n, o, p* account for background fluorescence, initial cell protein concentration, and other factors not specific to the TU. By differentiating the equations above with respect to *C*, we see that *r*_*xr*_, *r*_*xy*_, *r*_*sr*_, *r* _*sy*_ are the relative expression rates of the RFP TU and the YFP TU with respect to the CFP TU in each of the two dynamical regimes,

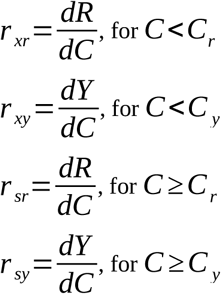

This piece-wise linear model was fitted to the phase space trajectories of all samples containing each plasmid (Figure 4A,B,D,E). Relative expression rates were fitted with standard error 6.53+/-7.00% (mean +/standard deviation). The dynamical regimes observed are thus characterized by their constant relative expression rates.

**Figure 4.**
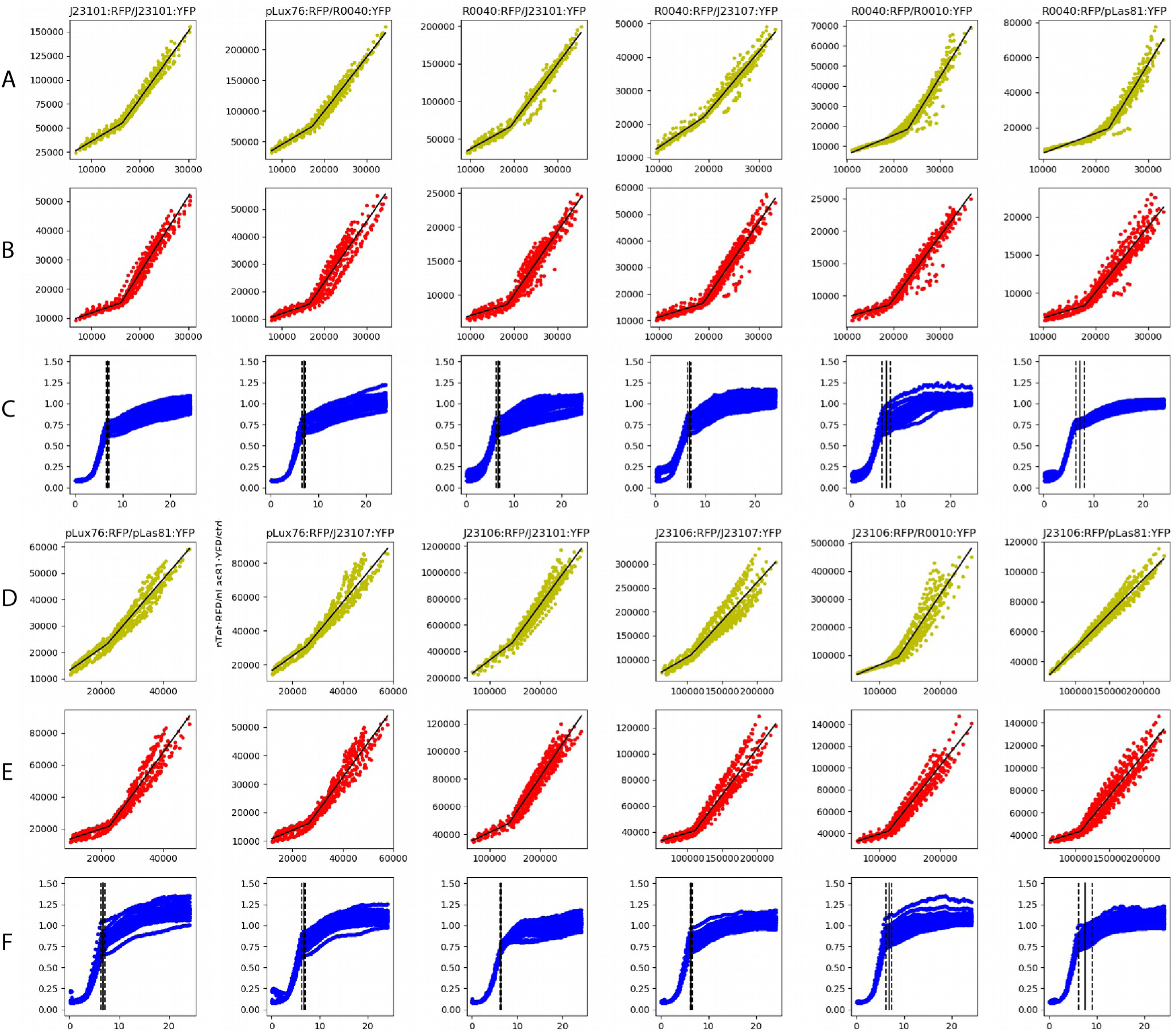
Phase plots with linear fits for YFP vs CFP (A,D), RFP vs CFP (B,E). Inflection points were mapped to time (see figure 3) and compared to OD traces (C,F). Black lines show mean and standard deviation (dashed lines) of times of inflections, clearly corresponding to the transition to stationary growth regime. All fluorescence measured in AU, time measured in hours.

## Dynamical regimes of plasmids correspond to changes in cellular context

To determine the relation between dynamical regime changes and dynamic changes in **cellular context** over the growth cycle, we determined the time at which they occur. We estimated the times *t* _0_ at which CFP fluorescence intensity *C* (*t*) =*C*_*c*_ and *C* (*t*) =*C*_*y*_ for phase space trajectories of each replicate for every plasmid (Figure 3AB). The resulting crossing times *t* _0_, which corresponded to inflection points in the fitted piecewise linear models, were mapped to growth by examining the optical density (OD) curve at the same times (Figure 3C). In Figure 4C,F we show the mean and standard deviation of the crossing times where the dynamical regime of TU expression changes showing that they clearly correspond to the transition to stationary growth regime. Hence TU expression rates change during the transition from exponential to stationary growth regimes, but the relative expression rate remains constant in each growth regime. The dynamical regimes observed in the phase space plots correspond to two distinct **cellular contexts** in which relative expression rates are constant.

Since their relative expression rates were constant in each cellular context, we can characterised the TU expression dynamics by a single parameter for each context. Equivalently, two parameters characterized the TU expression rates over a dynamically changing **cellular context**.

### A two-factor mathematical model can explain phase space trajectories

The host cell provides a range of factors that facilitate protein expression, and transcriptional units may be more or less sensitive to changes in these factors. Changes in **cellular contexts** are characterized by variation in many factors including RNA polymerase and ribosome levels, central metabolism, and sigma factors^37^. For fluorescent proteins these resources include those required for maturation. Many alternative models of differing complexity could explain the piecewise linear behavior of phase space trajectories. Here we propose a simple model with two main factors affecting TU expression rates.

Each factor *F*_*x*_ and *F*_*s*_ represents a basket of resources available in exponential and stationary growth regimes respectively. If we approximate that these resources are exclusively available in their corresponding **cellular contexts**, and propose a simple enzyme-substrate kinetic, the expression rate of a given TU can be modelled as,

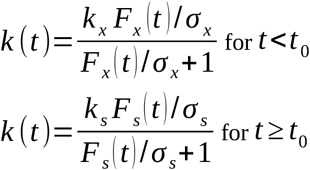

where *t*_0_ is the time of growth regime change, *k*_*x*_, *k*_*s*_ are the maximum expression rates, and *σ*_*x*_, *σ*_*s*_ are the sensitivities of the TU to each resource factor.

In resource limiting conditions *k* →*k*_*x*_*F*_*x*_ (*t*)/ *σ*_*x*_ in exponential growth regime, and *k* →*k*_*s*_*F*_*s*_ (*t*)/*σ*_*s*_ in stationary regime. By the same definition, the expression rate of the CFP TU is given by *k*_*c*_ → *k* _*xc*_ *F*_*x*_ (*t*) / *σ* _*xc*_ during exponential growth, and *k*_*c*_ →*k*_*sc*_*F*_*s*_ (*t*) / *σ* _*sc*_ during stationary growth. Hence the relative expression rate of a TU with respect to the reference CFP TU is given by,

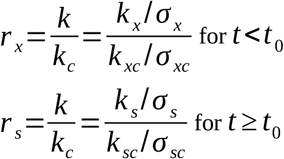

So that for resource limiting conditions, the potentially noisy time variation in resources is eliminated and the TU relative expression rates are constant in each **cellular context**, depending only on intrinsic properties of the TUs. Note that limiting resources is the case when all resources are actively involved in gene expression, primarily from the genome. Similarly for resource rich conditions (*F* →∞) we find that expression rates depend only on the intrinsic maximum rate of the TU. This result then suggests that relative sensitivities *σ*_*x*_ / *σ*_*xc*_, *σ*_*s*_ / *σ*_*sc*_ and/or relative maximum expression rates *k*_*x*_ / *k*_*xc*_, *k*_*s*_ / *k*_*sc*_ may change as cells transition between growth regimes or **cellular contexts**.

### TU relative expression rates are dependent on compositional context

For independent TUs their relative expression rates in each growth regime should not depend on adjacent TUs. That is, in any plasmid the relative expression of a given TU should be the same, irrespective of the adjacent TU relative expression rate. Competition for resources can be a significant factor in **compositional context** dependence of TUs. We measured the correlation between mean RFP and YFP relative expression rates in the same plasmid, and found no relation in exponential (Pearson correlation coefficient 0.115) nor stationary (Pearson correlation coefficient 0.208) growth regimes. This suggests that competition for resources was not a major factor in variation in relative expression rates of TUs.

Using our set of 12 plasmids we then compared the relative expression rates of the same TU with different adjacent TUs, or different **compositional contexts**. To examine the variance between contexts, we fitted the piece-wise linear model separately to every replicate of all plasmids (30 each, see Supplementary methods, figure S6-17 for full data). The combinations of RFP and YFP TU and their mean relative expression rates fitted to each replicate in each growth phase (*r*_*x*_ and *r*_*s*_) are shown in figure 6. Figure 5 shows the mean and standard deviation of these parameter estimates.

**Figure 5:**
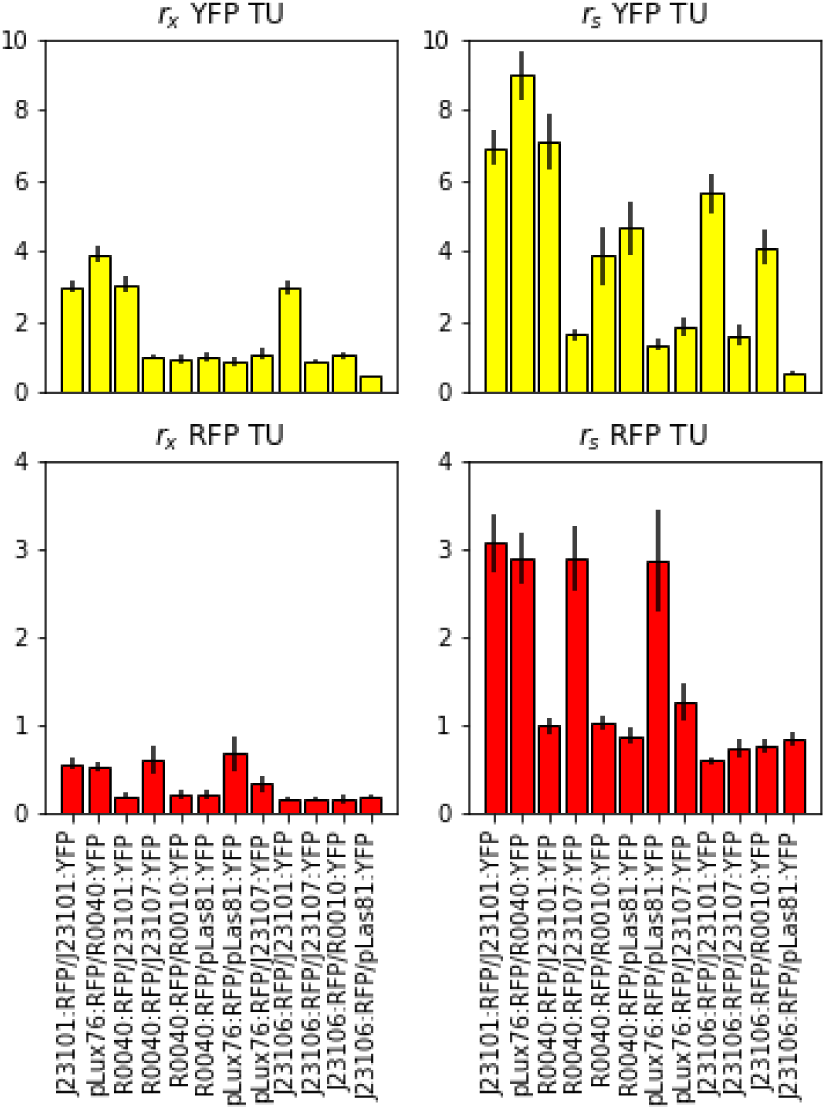
(Top row) Relative expression rates of YFP TUs in (left) exponential and (right) stationary regime. (Bottom row) Relative rates of RFP TUs in each growth regime. Error bars show one standard deviation of parameters fitted to each replicate separately.

**Figure 6.**
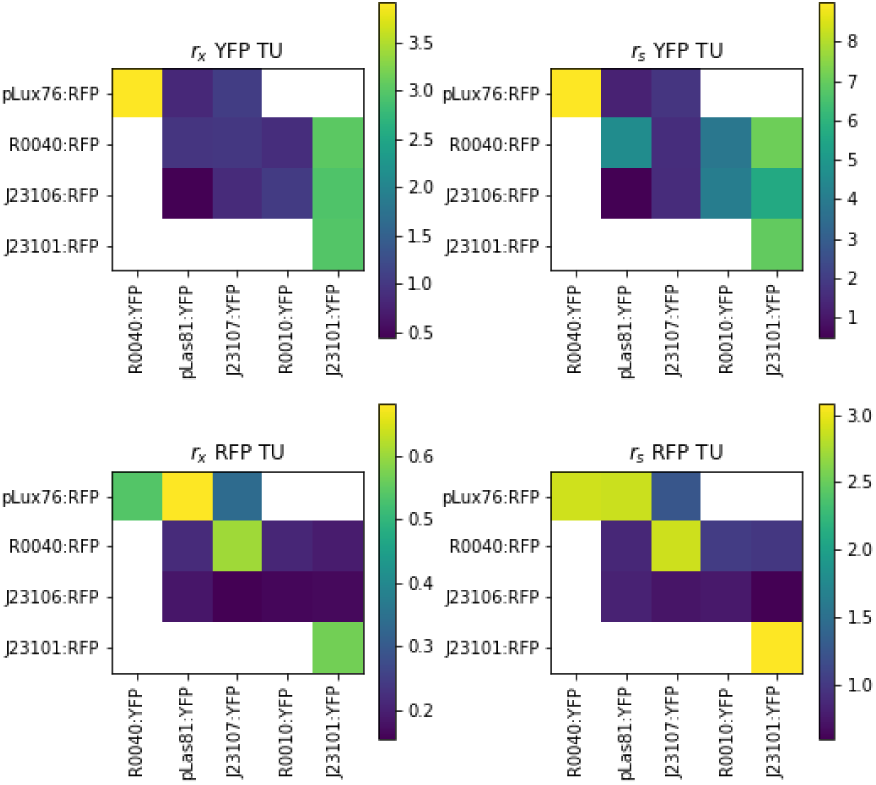
Combinations of TUs contained in the 12 plasmids considered in this study. (Top row) Relative expression rates of the YFP TU in exponential and stationary growth regimes (left, right) respectively. (Bottom row) Relative expression rates of the RFP TU during exponential and stationary growth (left, right). Values shown are for the mean parameters of piece-wise linear models fitted separately to all replicates for each plasmid.

To determine the effect of compositional context on these TUs, we performed a Kruskal-Wallis test^38^ on each TU grouped by **compositional context** (see table 1). The results show that all TUs were affected by compositional context in at least one of their relative expression rates. However we note that while significant, these effects were modest in most cases. We calculated pairwise fold-changes in mean relative expression rates between compositional contexts for each TU (figure 7). The results showed that 92% of context changes caused less than 3-fold change in relative expression rate, and 74% caused less than 2-fold change. We also note that the piecewise linear form of TU relative expression rates was maintained across **compositional contexts**. This means that in each combination of **cellular** and **compositional context** there was a different characteristic parameter of TU gene expression dynamics. For most TUs this characteristic parameter differed by less than 2-fold across compositional contexts.

**Table 1.** Kruskal-Wallis p-values for TU relative expression rates in exponential and stationary phase in different compositional contexts, significant differences (p<0.05) are marked in red.

| TU name | p-value, $r_x$ | p-value, $r_s$ | No. of contexts |
| --- | --- | --- | --- |
| pLux76:RFP | $1.96 \times 10^{-12}$ | $1.29 \times 10^{-13}$ | 3 |
| R0040:RFP | $8.03 \times 10^{-15}$ | $2.78 \times 10^{-18}$ | 4 |
| J23106:RFP | $5.38 \times 10^{-5}$ | $3.46 \times 10^{-14}$ | 4 |
| J23101:YFP | 0.579 | $7.11 \times 10^{-11}$ | 3 |
| R0010:YFP | $1.84 \times 10^{-4}$ | $7.60 \times 10^{-2}$ | 2 |
| J23107:YFP | $1.83 \times 10^{-9}$ | $3.60 \times 10^{-3}$ | 3 |
| pLas81:YFP | $5.43 \times 10^{-15}$ | $6.59 \times 10^{-18}$ | 3 |

**Figure 7.**
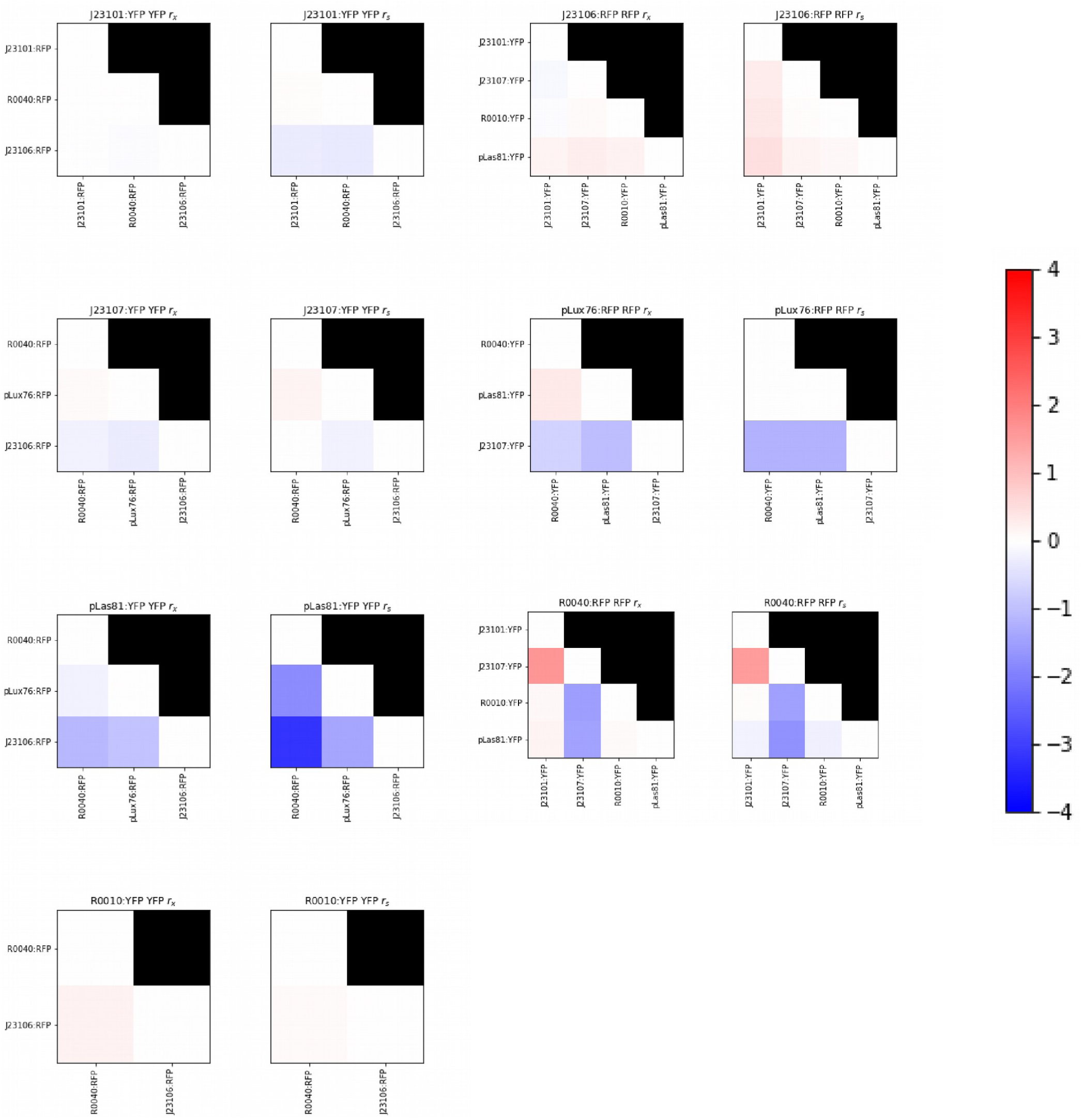
Log base 2 fold changes in mean relative expression rates (exponential and stationary phase) for each TU present in more than one compositional context. Each heatmap shows the fold change for pairwise comparisons between TU relative expression rates in the given compositional contexts (rows and columns of each heatmap).

### Accounting for cellular and compositional contexts in gene circuit design

We characterized gene circuits from the trajectories of 12 multi-fluorescent plasmids through a phase space parameterized by reporter expression levels. We considered the full culture growth cycle, and hence we observed dynamically changing **cellular context** which could be separated into two distinct linear dynamical regimes. In each dynamical regime the relative expression rates of TUs were constant. Fitting a piecewise linear model and mapping the inflection point of phase trajectories to time showed that the dynamical regimes corresponded closely to exponential and stationary growth regimes. Therefore the relative TU expression rates changed during the transition from exponential to stationary growth regime, but were constant during each regime or **cellular context**.

We present a two-factor mathematical model for cellular resources that accounts for this behaviour. This model is based on a simple transition of available resources for TU expression between **cellular contexts**. With resource levels that are mutually exclusive to their respective **cellular contexts** and present at rate-limiting or saturating levels this model predicted a piecewise linear dynamic as observed. In this model the characteristic slopes of the phase plots were due to differing relative sensitivities to resources, or to the ratio of maximum possible expression rates. Both of these quantities might be affected by the **compositional context** in which the TU is measured.

Comparing TU expression in different plasmids, we observed that relative expression rates were significantly affected by their neighbouring TUs, hence **compositional context** had an effect on circuit behaviour. While mostly less than 2-fold changes, these context effects could affect circuit functionality considerably, for example in the design of a balanced toggle switch. However, piecewise linear dynamics were observed for TUs in all **compositional** and **cellular contexts**. Hence the dynamics of TUs in a given context was dependent on a single unknown parameter, the relative expression rate with respect the reference TU.

Our results give the following equation to predict the expression rate of a TU in a gene circuit:

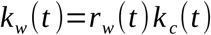

where *k*_*c, w*_ (*t*) is the expression rate of the reference TU in the context of interest *w*. For bulk culture 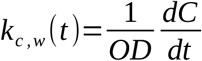, where *C* is the expression level of the reference TU in the given context. For constitutive TUs we showed that *r*_*w*_ (*t*) was a constant in both exponential and stationary growth regimes, but varied with compositional context. Hence the context *w* must incorporate both **compositional** and **cellular context**.

For TUs regulated by either chemical inducers or other TUs, this characteristic slope will be given by a time-varying regulatory function. Consider the case of an RFP TU repressed by a protein *P*. A simple Hill function model of the regulatory function would be,

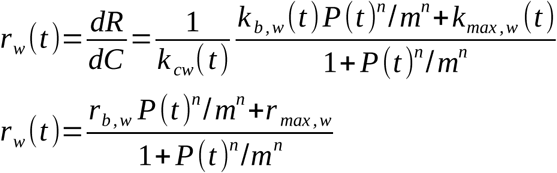

where *m, n* are constants. The functions *k*_*b, w*_, *k*_*max, w*_ are the basal and maximal expression rates, and *r*_*b, w*_, *r* _*max, w*_ are the basal and maximal relative expression rates in the context *w*.

Since repressed TUs followed piecewise linear dynamics in our experiments *r*_*b, w*_, *r* _*max, w*_ would be constant in a given context. For fixed driving protein concentrations, *r*_*w*_ would also be constant leading to a time-invariant regulatory function. Computing the slope of the phase space trajectory hence allows estimation of the parameters of regulatory functions that is independent of time variation in expression rates.

## Summary

This work highlights the importance of accounting for context in the characterization of gene circuits. Our approach provides a framework for design of gene circuits composed of TUs. A common standard reference TU may be chosen and measured in the cellular contexts of interest. Each TU in the design toolkit would be characterized by its phase space slopes in each context. During exponential and stationary growth regimes, the dynamics of each TU in any circuit may then be predicted by the output of the reference TU scaled by a single characteristic parameter, and regulatory functions can be expressed in terms of this characteristic. For other dynamically changing **cellular contexts** more complex models may be required, which could similarly be parameterized from the phase space trajectory slopes. It remains for future studies to characterize a broader range of such models.

Our results also show that **compositional context** within gene circuits significantly affected component TU expression rates, but mostly less than 3-fold variation (92%). Competition for resources between TUs was not a major factor in this variation. We also found that the functional form of variation in relative expression rates with respect to the reference TU was unaffected by changes in **compositional context**. **Compositional context** arises from many complex mechanisms making it unlikely that simple models can capture their effects. This means that TUs should be characterized in their **compositional context** of interest, or free parameters that account for their effects must be incorporated into models. In our experiments this would require an unknown scaling constant in each discrete **cellular context**. Since the compositional context effects appear to be limited in scale, their effects could be assessed by simulating gene circuits with a range of values for these free parameters.

Our approach of phase space characterization will thus aid in genetic circuit design by providing simple models with few parameters to incorporate into the design-build-test cycle. As well as enabling fitting to data, these models will allow more accurate simulation^8^ of gene circuit dynamics for parameter sweeps, for example to assess the probability of circuit design success. They will also allow the definition of contextual constraints on circuit operation^1^. The use of a common standard reference TU will enable comparable and sharable characterization of gene circuits and their component TUs. This dynamical characterization will therefore enable more reliable design of synthetic gene circuits as well as understanding of naturally occurring genetic systems.

## Methods

### Strains

For plasmid selection and stock, the strain used was *Escherichia coli* TOP10 (Invitrogen). For making the growth assays the strain used was *Escherichia coli* MG1655Z1 malE, which has constitutive levels of LacI and TetR repressors^39^ (Addgene).

### Modular system for DNA assembly

### Golden Gate Cloning

Transcriptional Units (TU) were built using level 0 DNA parts, consisting of seven Promoters, two Ribosome Binding Sites (RBS), two Coding Sequences (CDS), two terminators and two acceptor plasmids (see details in Supplementary methods). They were assembled in order using Golden Gate Cloning, obtaining a level 1 TU^40,41^. For each Golden Gate reaction 1µl of T4 DNA Ligase Buffer 10X (NEB), 1µl of T4 DNA Ligase 20U/µl (NEB) and 1 µl of BsaI 10 U/µl (NEB) were added to 7 µl of a mixture of the DNA parts used for each TU, with a final volume of 10 µl in 0,2 ml tubes. The thermocycler program used was 2 minutes at 37°C and 3 minutes at 16°C during 20 cycles, then 5 minutes at 50°C and 10 minutes at 80°C (for details see the Supplementary methods).

### Gibson Assembly

For building the three fluorescent reporters plasmids, a destination vector was built first using Gibson Assembly, named 1X_p15a_Cyan. For this, linear DNA pieces were obtained through PCR with Phusion Polymerase (Invitrogen) from template plasmids (for details see the Supplementary material). For one Gibson reaction 4.5 µl of Gibson Master Mix were added to 1.5 µl of a mixture of the DNA pieces in a 0,2 ml tubes and incubated at 50°C in a thermocycler for one hour^42^.

For building the three fluorescent reporters plasmids, each TU was amplified by PCR in the same way as for the destination vector and were introduced in different combinations in the destination vector using Gibson Assembly, but in this case with the help of Unique Nucleotide Sequences (UNSes^43^). In each plasmid there was a TU containing RFP and UNSes U1 and U2, and other containing YFP and UNSes U2 and UX, and the destination vector had the UNSes U1 and UX, which led to an ordered assembly, obtaining 12 plasmids (for details see the Supplementary methods).

### Primer design

The following features were considered for primers design: length less than 60 base pairs; Tm ≤ 60°C; less than four C or G at the ends, specially at the 3’ end and 45% ≤ CG% ≤ 60%.

### DNA pieces purification from agarose gel

The linear pieces obtained from PCR reactions for Gibson Assembly were run in 2% w/v agarose gel. The bands were purified using the Wizard® Plus SV Gel and PCR Clean-Up System kit (Promega), using the Quick Protocol specified by the manufacturer. The quality and concentration of the parts were quantified using the Synergy HTX plate reader (BioTek) and Gen5 software with the Take3 plate (BioTek). The pieces were stored at −20°C.

### Bacteria transformation

For plasmid cloning, CCMB80 chemocompetent *Escherichia coli* TOP10 were prepared and transformed. For Gibson Assembly the complete content of the reaction was added to the cells (i.e. 6 µl), while for Golden Gate Cloning only 5 µl were added. The cells were then subjected to a heat shock at 42°C for 1 minute in a thermoregulated bath. Then 250 µl of liquid LB media were added to the cells and incubated at 37°C for one hour. In solid LB media with antibiotic, 100 µl of the cells were plated and incubated overnight at 37°C. For growth assays, the same protocol was used with MG1655Z1 malE cells with plasmids already verified, except that in this case 50 µl were plated in solid LB media with antibiotic.

### Selection and storage of positive colonies

Two to three colonies of TOP10 transformed cells were selected to verify through PCR with GoTaq DNA Polymerase (Promega) if the plasmids they contained were the desired constructions. Primers for specific regions of the plasmids were used (for details see the Supplementary methods). The PCR products were run in 2% w/v agarose gels. Once a colony was verified to contain the correct plasmid, a liquid culture in LB media was left growing overnight. From this culture, 500 µl were used for storage at −80°C, adding 500 µl of 50% v/v glycerol in a cryotube. For MG1655Z1 malE cells, liquid cultures were left growing and stored the same way (for details see the Supplementary methods).

### Plasmid extraction and purification

Liquid cultures of the transformed cells were used for plasmid extraction and purification using the Wizard® Plus SV Minipreps DNA Purification System kit (Promega). The plasmids quality and concentration were quantified using the Synergy HTX plate reader (BioTek) and Gen5 software (BioTek). The plasmids were stored at −20°C.

### Growth assays

For growth assays M9 media was used with 0.4% w/v Glucose and 0.2% casamino acids. Optical density of the colonies that contained the different plasmids and fluorescence of RFP, YFP and CFP were measured for 24 hours, every 15 minutes, at 37°C with constant shaking in 96 well black plates (Thermo). Each growth assay contained 10 replicates and was repeated on 3 different days. The measurements were taken with a Synergy HTX plate reader (BioTek) with Gen5 software (BioTek) (for details see the Supplementary methods).

### Data analysis

Data from growth assays were stored in a custom database (data files and code available at github.com/SynBioUC/flapjack). The analysis was performed using Python and packages Numpy^44^, Scipy^45^, Matplotlib^46^, Pandas^47^, SQLAlchemy (www.sqlalchemy.org), and Jupyter (jupyter.org). Piecewise linear models were fitted using the Scipy *curve_fit* function, which implements the Trust Region Reflective algorithm. To avoid initial noisy data points and focus on the period at which the inflection point occurred, as well as remove data bias from long stationary phase, we fitted to data from time point 20 (approximately 5 hours) to time point 40 (approximately 10 hours). Stated standard deviations of parameters from complete data were those returned in the covariance matrix by the *curve_fit* function. Figure S6-17 shows all the phase space trajectories and linear model fits for each replicate of each plasmid. Standard deviations stated were the population standard deviation of all replicates. To determine the influence of compositional context upon *r*_*x*_ and *r*_*s*_ variation, a Kruskal-Wallis test^38^ was performed for each TU, grouped by plasmid, using the scipy function *kruskal*.

## AUTHOR INFORMATION

### Author Contributions

All authors have given approval to the final version of the manuscript. MAMS performed experiments and analyzed data. IÑ and TM performed experiments. TJR, GAVP, CAR and AV analyzed data. TJR, MAMS, and FF designed the study and wrote the manuscript.

### Funding Sources

TJR and MAMS were supported by FONDECYT Iniciacion No. 11161046. FF was funded by Instituto Milenio iBio – Iniciativa Científica Milenio MINECON.

## ACKNOWLEDGMENT

The authors would like to thank Guillermo Yañez for his help uploading to SynBioHub.

