## Supplementary methods for "Phase space characterization for gene circuit design"

### Methods

#### Strains

For plasmids selection and stock, the strain used was *Escherichia coli* TOP10 One Shot™ chemocompetents (Invitrogen) [1]. For making the growth assays the strain used was *Escherichia coli* MG1655Z1 malE, which has constitutive levels of LacI and TetR repressors [2].

### Modular system for DNA assembly

#### Golden Gate Cloning

Transcriptional Units (TU) were built using level 0 DNA parts or modules, consisting of seven Promoters, two Ribosome Binding Sites (RBS), two Coding Sequences (CDS), two terminators and two acceptor plasmids (*CIDAR MoClo*, Addgene Kit #1000000059). The names of the parts and combinations are detailed in table S1 and S2.

The parts sequences were contained in level 0 plasmids and were flanked by a restriction site for BsaI, which was GAGACC/CTCTGG, and a four-nucleotide small sequence named from A to E, according to the CIDAR MoClo system based on Golden Gate Cloning[3]. Promoters were flanked by sequences A and B, RBSs were flanked by sequences B and C, the CDSs were flanked by C and D, the terminator were flanked by D and E, and the acceptor plasmids contained sequences A and E. This allowed the parts to be assembled in order and the restriction sites were deleted, obtaining a level 1 TU [3,4]. Sequences A to E are detailed in table S3.

For reactions, a mix of the level 0 modules was made, calculating equimolar mounts of the modules obtaining 10 fmol of each part in a volume of 10 µl. From this mixture, 7 µl were used for a Golden Gate reaction, and 1µl of T4 DNA Ligase Buffer 10X (NEB), 1µl of T4 DNA Ligase 10U/µl (NEB) and 1 µl of BsaI 20 U/µl (NEB) were added to the mixture for each TU, with a final volume of 10 µl in 0.2 ml tubes. The thermocycler program used was 2 minutes at 37°C and 3 minutes at 16°C during 20 cycles, then 5 minutes at 50°C and 10 minutes at 80°C.

Figure S1 shows the calculations for the mix of level 0 parts.

**Table S1. Transcriptional Units (TU) that contain RFP**

| Promoters | RBS | CDS | Terminator | Acceptor plasmid |
|---|---|---|---|---|
| A_J23101_B | B_BCD2_C | C_RFP_D | D_ECK0818_E | 12X |
| A_J23106_B | B_BCD2_C | C_RFP_D | D_ECK0818_E | 12X |
| A_R0040_B | B_BCD2_C | C_RFP_D | D_ECK0818_E | 12X |
| A_pLux76_B | B_BCD2_C | C_RFP_D | D_ECK0818_E | 12X |

**Table S2. Transcriptional Units (TU) that contain YFP**

| Promoters | RBS | CDS | Terminator | Acceptor plasmid |
|---|---|---|---|---|
| A_J23101_B | B_BCD12_C | C_YFP_D | D_ECK9600_E | 23X |
| A_J23107_B | B_BCD12_C | C_YFP_D | D_ECK9600_E | 23X |
| A_R0040_B | B_BCD12_C | C_YFP_D | D_ECK9600_E | 23X |
| A_R0010_B | B_BCD12_C | C_YFP_D | D_ECK9600_E | 23X |
| A_pLas81_B | B_BCD12_C | C_YFP_D | D_ECK9600_E | 23X |

**Table S3. Four-nucleotide flanking sequences for Golden Gate CIDAR Moclo**

| Module | Sequence name | Sequence |
|---|---|---|
| Promoter | A | GGAG |
| | B | TACT |
| RBS | B | TACT |
| | C | AATG |
| CDS | C | AATG |
| | D | AGGT |
| Terminator | D | AGGT |
| | E | GCTT |

**A** $$Module\ fmoles = \left(\frac{[{}^{ng}/_{\mu l}]}{Module\ length\ (bp)}\right) * \left(\frac{10^6}{660}\right)$$

**B** $$Volume\ relation = \frac{needed\ mount\ of\ fmoles}{module\ fmoles}$$

**C** $$Module\ volume\ to\ add = \left(\frac{final\ mix\ volume}{\begin{matrix} addition\ of\ volume\ relation \\ of\ the\ modules \end{matrix}}\right) * \begin{matrix} vol.relation\ of \\ one\ module \end{matrix}$$

**Figure S1. Calculations of equimolar quantities mixture of level 0 parts for Golden Gate Cloning.** A) Formula for determining the femtomole concentration of each module from its concentration in nanograms per microlitre. The module length is the plasmid size in base-pairs, $10^6$ is a conversion factor for femtograms and 660 corresponds to the average molecular weight of a pair of nucleotides. B) To determine the volume to add of each part, first is necessary to calculate the relation between the parts. For this, the volume relation of a part is calculated dividing the needed mount of femtomoles (10fmol) in the mount of femtomoles determined in A) for each part or module. C) In this formula, the final mix volume corresponds to the final volume of the mixture of parts (10 μl), which is divided by the addition of all the volume relations of the parts to assemble, and this value is multiplied by the volume relation of the part to add. This formula gives the volume of that part that is needed to add to the final mixture of parts.

### Gibson Assembly

For building the three TUs plasmids, a destination plasmid was built first using Gibson Assembly. This plasmid was named 1X_p15a_Cyan. Gibson Assembly uses three enzymes for its reaction, allowing to assemble DNA molecules or pieces of big size. The reaction happens in 0.2 ml tubes at 50°C, and the enzymes used are a T5 5'-exonuclease (Epicentre), a high-fidelity polymerase, which is Phusion polymerase (Invitrogen) and a DNA Taq Ligase (NEB). In figure S2, the Gibson Assembly is described[5].

In order to assemble the TUs in the destination plasmid, linear DNA pieces were obtained through PCR with Phusion Polymerase (Invitrogen) from template plasmids, these templates are specified in table S4. The PCR mix is detailed in table S5 and the program used was 98°C for 30 seconds, then 35 cycles with 98°C for 10 seconds, 65°C for 30 seconds and 72°C for 1 to 3 minutes depending on the amplified piece; and finally, 72°C for 7 minutes.

The PCR products were run on agarose gels, the concentration was 2%w/v in TAE buffer 1X for pieces smaller than 3kb and 1%w/v in TAE buffer 1X for pieces bigger than the same size. All the gels were run at 100V for 30-45 minutes. The bands were purified from the gel and stored as specified in the "DNA pieces purification from agarose gel" section in this file.

For the Gibson reaction, a mix of linear DNA parts was used. To determine how much of each piece was needed, their equimolar quantities in picomoles had to be calculated for a final volume of 4 µl for this mixture, figure S3 shows the formulas for these calculations. For one Gibson reaction, 4.5 µl of Gibson Master Mix were added to 1.5 µl of the mixture of the DNA pieces in a 0.2 ml tubes and incubated in a thermocycler at 50°C for 5 minutes and then 60 minutes at the same temperature[5]. Table S6 shows the Gibson mix composition used for the reaction.

For building the three TUs plasmids, each TU was amplified by PCR in the same way as for the destination plasmid and were introduced in different combinations in the destination vector using Gibson Assembly, but in this case with the help of Unique Nucleotide Sequences[6] (UNSes), which were flanking the TUs inside their plasmids. The TUs containing RFP had UNS1 and UNS2 sequences, while the TUs containing YFP had UNS2 and UNSX sequences and the destination plasmid had the UNS1 and UNSX sequences, allowing an ordered assembly. The primers used for amplifying the TUs and the destination plasmid are detailed in Table S7. The UNSes used are described in Torella *et al.*, 2014[6].

This way, 12 plasmids of three TUs, were obtained, the combination of TUs are specified in table S8, and the TUs are named by the promoter contained.

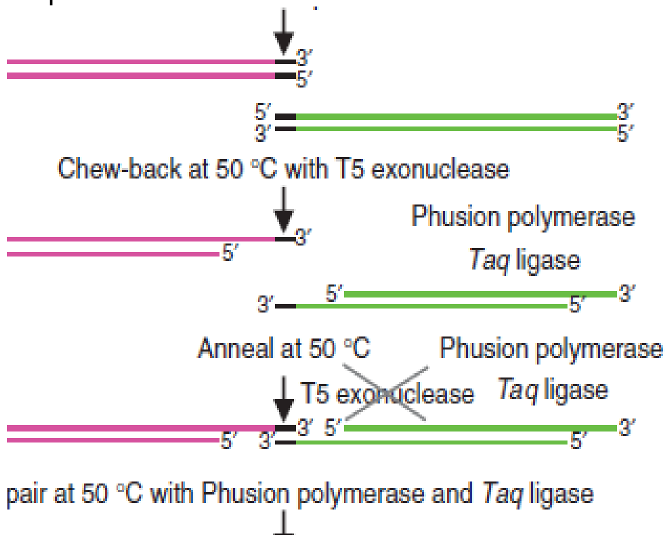


**Figure S2. Gibson Assembly reaction scheme.** All the steps of the reaction occur in the same tube and at the same temperature. First, the T5 5'-Exonuclease cleaves the 5'- end of the linear DNA molecule, and these join by complementarity and the Polymerase fills the spaces between molecules. Finally, the DNA Taq ligase fixes and joins the molecules (Figure from Gibson *et* al., 2009[5]).

**Table S4. Template plasmids and primers sequences for pieces amplification for 1X_p15a_Cyan destination plasmid**

| Piece | Primer | Primer Sequences | Template plasmid | Piece size (bp) |
|---|---|---|---|---|
| A | U1F<br>UXR | CATTACTCGCATCCATTCTCAGGCTG<br>GGTGGAAGGGCTCGGAGTTGTGG | pDestBAC | 115 |
| B | 1X_p15aF<br>pGreenR | GATACATAGATTACCACAACTCCGAGCCCTTCCACCT-ACTAGTAGCGGCCGCTGCAGTC<br>TGCGCCGGTTGCATTCGATTCCTG | T52V2 | 700 |
| C | pGreenF<br>1X_p15aR | CCTGAGCGAGACGAAATACGCGATC<br>GAGACGAGACAGCCTGAGAATGGATGCGAGTAATGTC-TAGAAGCGGCCGCGAATTCCAG | STDSTD7 | 3105 |

**Table S5. PCR reaction mix for Phusion Polymerase**

| Reagent | Mix 1X (μl) | 1X_p15a_Cyan (3,1X) |
|---|---|---|
| 5X Buffer | 6 | 18.6 |
| $H_2O$ | 20.6 | 63.86 |
| dNTPs | 0.6 | 1.86 |
| Primer forward | 0.75 | 2.325 |
| Primer reverse | 0.75 | 2.325 |
| Phusion | 0.3 | 0.93 |
| Template DNA | 1 | - |
| Final volume | 30 | - |

One reaction was made for each piece of DNA. One microlitre of template DNA was added to 29 μl of the reaction mix. Three reactions were made for building the destination plasmid, and there 0.1μl is added for accounting for pipetting errors.

**A** $$Piece\ pmoles = \left( \frac{[^{ng}/_{\mu l}] * 1000}{Piece\ length\ (bp) * 650} \right)$$

**B** $$Volume\ relation = \frac{Factor}{piece\ pmoles}$$

**C** $$Piece\ volume\ to\ add = \left( \frac{final\ mix\ volume}{\substack{addition\ of\ volume\ relation \\ of\ the\ pieces}} \right) * \substack{vol.relation\ of \\ one\ piece}$$

**Figura S3. Calculations of equimolar quantities mixture for DNA pieces for Gibson Assembly.** A) Formula for calculating the picomoles of each piece from its concentration in nanograms per microlitre. To convert nanograms to picograms, a factor of 1000 is used, the piece length corresponds to the piece size in base-pairs, and 650 is the average weight of a base-pair. B) To determine the volume to add to the mixture of pieces, first the volume relation of each piece is calculated by dividing a factor in the piece picomoles. The factor used could be 0.01 for pieces greater than 2500 bp, 0.03 if the piece size is from 200 bp and 2500 bp or 0.05 if the piece is smaller than 200 bp. C) Formula for determining the volume to add of one piece of DNA. The final volume corresponds to 4 µl, which is divided by the addition of all the volume relations of the pieces to assemble, and this is multiplied by the volume relation of the piece to add.

**Table S6. Reaction mix for Gibson Assembly**

| Reagent | Volume (µl) |
|---|---|
| Isothermal reaction 5X Buffer | 100 |
| T5 exonuclease 10U/ µl | 2 |
| Phusion Polymerase 2U/ µl | 6.25 |
| DNA Taq ligase 40 U/ µl | 50 |
| $H_2O$ | 216.75 |
| Final volume | 375 |

This mix can be stored in 15 µl aliquots up for a year[5], only 4.5 µl will be needed for one reaction.

**Table S7. UNSes sequences and primers for TUs and destination plasmid amplification**

| Template pieces | UNSes | UNSes Sequences | Primers | Primer sequence |
|---|---|---|---|---|
| 1X_p15a_Cyan | U1 | CATTACTCGCATCCATTCTCAGGCTG-TCTCGTCTCGTCTC | U1R | GAGACGAGACGAGACAG-CCTGAG |
| | UX | CCAGGATACATAGATTACCACAACTC-CGAGCCCTTCCACC | UXF | CCAGGATACATAGATTA-CCACAACTCCG |
| TU-RFP | U1 | CATTACTCGCATCCATTCTCAGGCTG-TCTCGTCTCGTCTC | U1F | CATTACTCGCATCCATT-CTCAGGCTG |
| | U2 | GCTGGGAGTTCGTAGACGGAAACAAA-CGCAGAATCCAAGC | U2R | GCTTGGATTCTGCGTTTG-TTTCCGTC |
| TU-YFP | U2 | GCTGGGAGTTCGTAGACGGAAACAAA-CGCAGAATCCAAGC | U2F | GCTGGGAGTTCGTAGAC-GGAAACAAAC |
| | UX | CCAGGATACATAGATTACCACAACTC-CGAGCCCTTCCACC | UXR | GGTGGAAGGGCTCGGAGT-TGTGG |

The UNSes sequences used are the ones described in Torella *et al.*, 2014[6].

**Table S8. TUs combinations introduces in the destination plasmid**

| | J23101 (YFP) | J23107 (YFP) | R0040 (YFP) | R0010 (YFP) | pLas81 (YFP) |
|---|---|---|---|---|---|
| J23101(RFP) | J23101/J23101 | - | - | - | - |
| J23106(RFP) | J23106/J23101 | J23106/J23107 | - | J23106/R0010 | J23106/pLas81 |
| R0040(RFP) | R0040/J23101 | R0040/J23107 | - | R0040/R0010 | R0040/pLas81 |
| pLux76 (RFP) | - | pLux76/J23107 | pLux76/R0040 | - | pLux76/pLas81 |

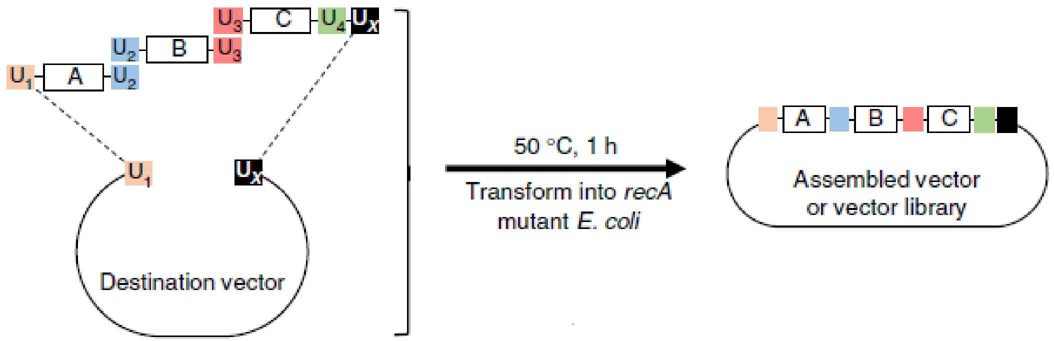


**Figure S4. Gibson Assembly using UNSes.** Each piece will be flanked by two UNSes sequences, which will be digested by the T5 5' Exonuclease, leaving complementary ends, so the last UNS sequence from one piece is the first sequence from the next piece of DNA. This allows the ordered assembly of the pieces[5,6] (figure modified from Torella et al., 2014[6]).

### Primer design

The following features were considered for primers design: length less than 60 base pairs; Tm ≤ 60°C; less than four C or G at the ends, specially at the 3' end and 45% ≤ CG% ≤ 60%.

### DNA pieces purification from agarose gel

The linear pieces obtained from PCR reactions for Gibson Assembly were run in 2% w/v agarose gel. The bands were purified using the Wizard® Plus SV Gel and PCR Clean-Up System kit (Promega), using the Quick Protocol specified by the manufacturer. The quality and concentration of the parts were quantified using the Synergy HTX plate reader (BioTek) and Gen5 software with the Take3 plate (BioTek). The pieces were stored at -20°C.

### Bacteria transformation

For plasmid selection, *Escherichia coli* TOP10 One Shot™ chemocompetent cells (Invitrogen) were transformed. For Gibson Assembly the complete content of the reaction was added to the cells, while for Golden Gate Cloning only 5 µl were added. The cells were then subjected to a heat shock at 42°C for 1 minute in a thermoregulated bath. Then 250 µl of liquid LB media were added to the cells and incubated at 37°C for one hour. In solid LB media with antibiotic, 100 µl of the cells were plated and incubated overnight at 37°C. For growth assays, the same protocol was used with MG1655Z1 malE cells with plasmids already verified, except that in this case 50 µl were plated in solid LB media with antibiotic.

### Selection and storage of positive colonies

Two to three colonies of TOP10 transformed cells were selected to verify through PCR with Gotaq Polymerase (Promega) if the plasmids they contained were the desired constructions. Primers for specific regions of the plasmids were used, table S9 shows the primers sequences in detail and table S10 shows the Gotaq mix composition. The primers used were for U1 and U2 sequences and for the terminators used (ECK0818 and ECK9600), to check the correct assembly of TUs. To check the correct assembly of the final plasmids, the same primers were used for the RFP and YFP TUs, and for the CFP TU, a primer for the J23101 promoter was used and a primer that anneals to the final end of CFP and its terminator. The PCR products were run in 2% w/v agarose gels and run at 100V for 30-45 minutes. Once a colony was verified to contain the correct plasmid, a liquid culture in LB media was left growing overnight. From this culture, 500 µl were used for storage at -80°C, adding 500 µl of 50% v/v glycerol in a cryotube (Invitrogen). For MG1655Z1 malE cells, liquid cultures were left growing and stored the same way.

**Table S9. Primers sequences for colony verification**

| TU | Primers | Primer sequence |
|---|---|---|
| CFP | STDF | TTTACAGCTAGCTCAGTCCTAGGTATTATGC |
| | CFPTR | GGTGGGCCTTTCTGCGTTTATAGCTTAGAGACC |
| RFP | U1F | CATTACTCGCATCCATTCTCAGGCTG |
| | 0818R | GGGCGATGGCCCACTACGTGGGTCTCTAAGCCCAA AAGTAAAAACCCGCCGAAGCGG |
| YFP | U2F | GCTGGGAGTTCGTAGACGGAAACAAAC |
| | ECK9600R | GCGATGGCCCACTACGTGGGTCTCTAAGCTTGAGA AGAGAAAAGAAAACCGCCGA |

**Table S10. Gotaq Polymerase reaction mix**

| Reagent | 1X (µL) |
|---|---|
| Green Buffer Gotaq 5X | 2 |
| $H_2O$ | 4.45 |
| $MgCl_2$ 25Mm | 1.6 |
| DNTPs | 0.2 |
| Primer forward | 0.25 |
| Primer reverse | 0.25 |
| Enzima Gotaq | 0.25 |
| Templado | 1 |
| Final volume | 10 |

The program used in the thermocycler was 95°C for 10 minutes, then 35 ciclos with 95°C for 30 seconds, 55°- 63°C for 30 seconds (depending on the pair of primers used), 72°C for 1:30 minutes. Finally, 72°C for 10 minutes.

#### Plasmid extraction and purification

Liquid cultures of the transformed cells were used for plasmid extraction and purification using the Wizard® Plus SV Minipreps DNA Purification System kit (Promega). The plasmids quality and concentration were quantified using the Synergy HTX plate reader (BioTek) and Gen5 software (BioTek). The plasmids were stored at -20°C.

#### Growth assays

For growth assays M9 media was used with 0.4% w/v Glucose and 0.2% w/v casaminoacids. Optical density of the colonies that contained the different plasmids and fluorescence of RFP, YFP and CFP were measured for 24 hours, every 15 minutes, at 37°C with constant shaking in 96 well black plates (Thermo). Each growth assay contained 10 replicates of each plasmid and was repeated on 3 different days, it included four wells of non-transformed bacteria and four wells of the M9 media used without bacteria as controls. Figure S5 gives an examplo of the colonies distribution. The measurements were taken with a Synergy HTX plate reader (BioTek) with Gen5 software (BioTek) (for details see the supplementary material methods). The CFP TU was used as a reference following the work of Kelly *et al*., 2009[7] and Rudge *et al*., 2016[8]. The M9 media composition is detailes in table S11.

To make the assays liquid cultures of the colonies were left growing over night (or 14-15 hours as a top) in M9 media supplemented with 0.4% w/v Glucose and 0.2% w/v casamino acids. The next day 25 ml of fresh M9 media was prepared as specified in table S11. This was distributed in 2 ml tubes, adding 1996 µl of media, and then 2 µl of kanamycin and 2 µl of the bacteria liquid culture was added to each tube to reach a final volume of 2 ml. Each colony was prepared in a separated tube.

Then 200 µl of each of the tubes was added to 10 wells for the colonies with the plasmids (10 replicates), and to four wells for each control. Then the plate was closed and sealed with parafilm and was carried to the plate reader.

<table>
<tr><th></th><th>1</th><th>2</th><th>3</th><th>4</th><th>5</th><th>6</th><th>7</th><th>8</th><th>9</th><th>10</th><th>11</th><th>12</th></tr>
<tr><td>A</td><td rowspan="4">M9 media without bacteria</td><td rowspan="4">M9 media with non transforme d bacteria</td><td colspan="5" rowspan="2">Colony with plasmid 1</td><td colspan="5" rowspan="2">Colony with plasmid 2</td></tr>
<tr><td>B</td></tr>
<tr><td>C</td><td colspan="5" rowspan="2">Colony with plasmid 3</td><td colspan="5" rowspan="2">Colony with plasmid 4</td></tr>
<tr><td>D</td></tr>
</table>

**Figure S5. Example of colonies distribution for growth assays.** Here half of the plate is shown. For each plasmid 10 replicates were used, and for each control 4 wells as indicated.

**Table S11. M9 media composition for growth assays**

| Reagent | Stock Solution concentration | Needed volume |
|---|---|---|
| Sterile destiled water | - | 14.45 ml |
| $MgSO_4 * 7H_2O$ | 1M | 50µl |
| M9 Salts* | 5X | 5 ml |
| Glucose | 20% v/v | 250 µl |
| $CaCl_2$ | 1M | 2.5 µl |
| Casaminoacids | 1% p/v | 5 ml |
| Final volume | - | 25ml |

*The M9 5X Salts are composed by $Na_2HPO_4$, $KH_2PO_4$, NaCl, $NH_4Cl$. The media components must be added in the shown order. The Glucose and casminoacids volumes used allow to obtain a final concentration of 0.4% y 0.2% respectively. M9 media must be prepared when is going to be used.

### Data analysis

Data from growth assays were stored in a custom database (database files and code in github.com/SynBioUC/flapjack). The analysis was performed using Python and packages Numpy[9], Scipy[10], Matplotlib[11], Pandas[12], SQLAlchemy, and Jupyter. Piecewise linear models were fitted using the Scipy *curve_fit*[10] function, which implements the Trust Region Reflective algorithm[10]. To avoid initial noisy data points and focus on the period at which the inflection point occurred, as well as remove data bias from long stationary phase, we fitted to data from time point 20 (approximately 5 hours) to time point 40 (approximately 10 hours). Figure S6-17 shows all the phase space trajectories and linear model fits for each replicate of each plasmid. Stated standard deviations of parameters were those returned in the covariance matrix. For model fitting, fluorescence time course data were first normalised by the maximum value and fitted with bounds [0,1] for parameters x,y of inflection point, and [0,2] for rx, rs.

To determine the influence of compositional context upon rx y rs variation, a Kruskal-Wallis test[13] was performed for each TU using the scipy function *kruskal*[10].

J23101:RFP/J23101:YFP/J23101:CFP

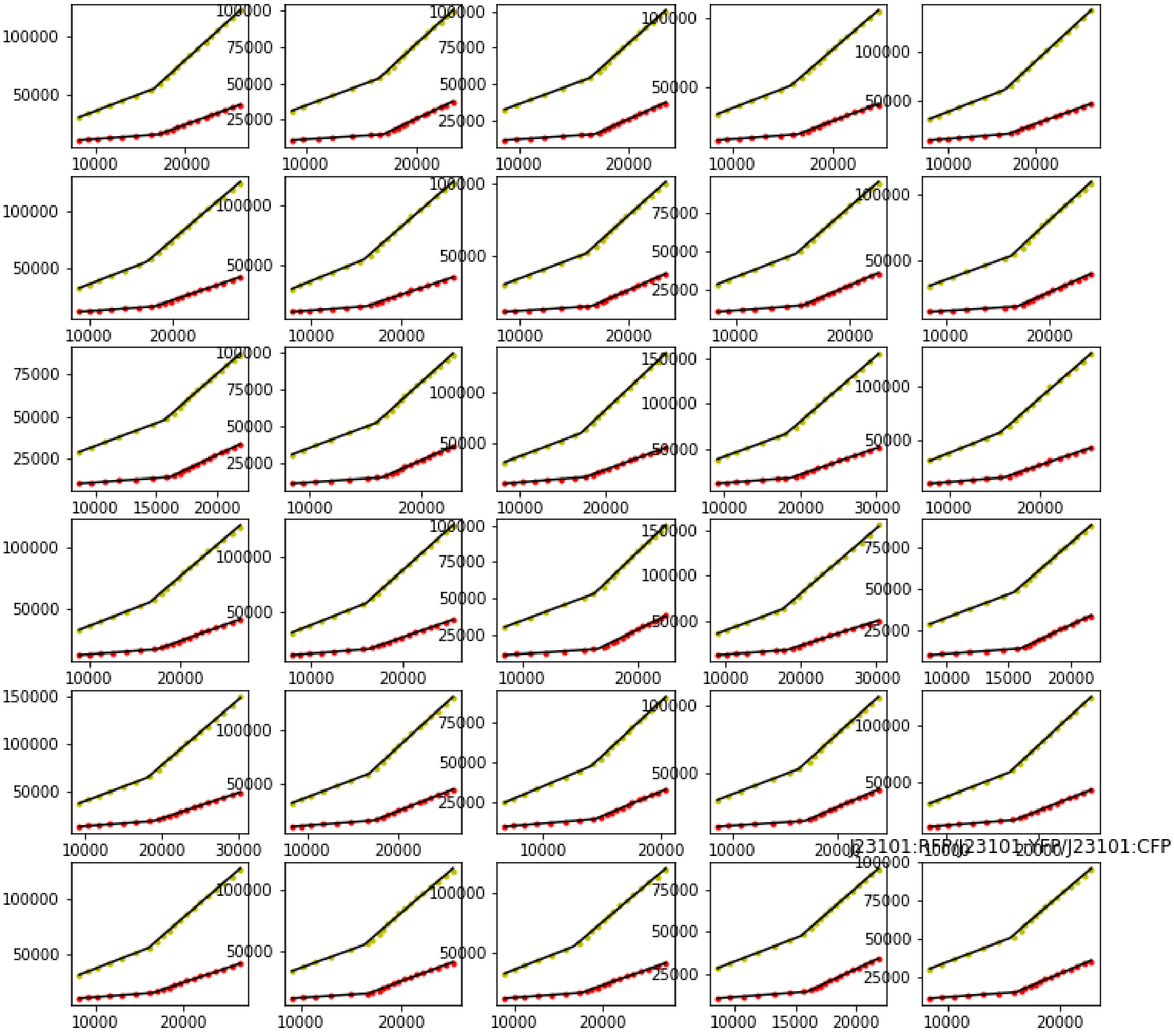

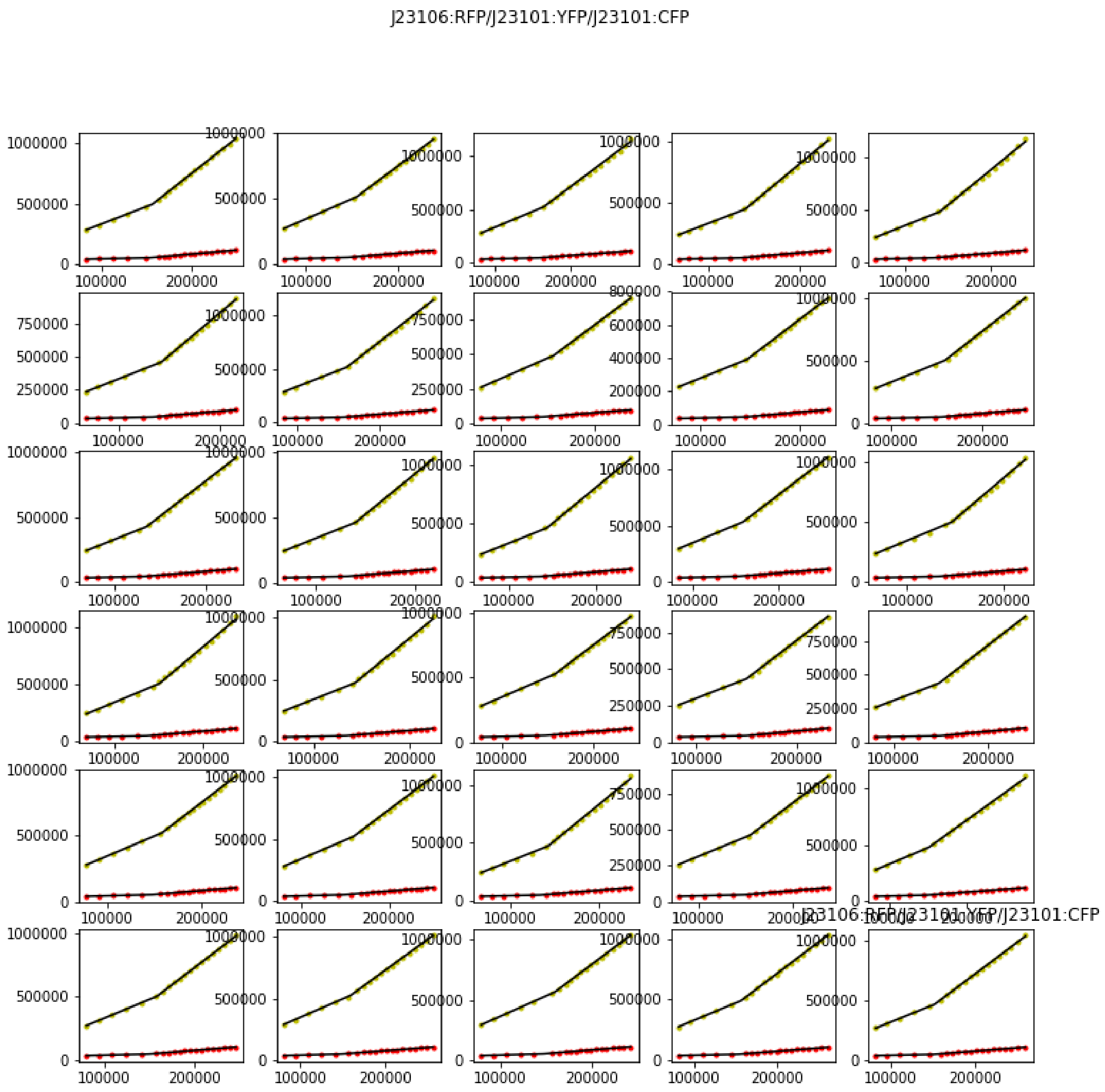


Figure S7. All phase space trajectories for plasmid J23106:RFP/J23101:YFP/J23101:CFP.

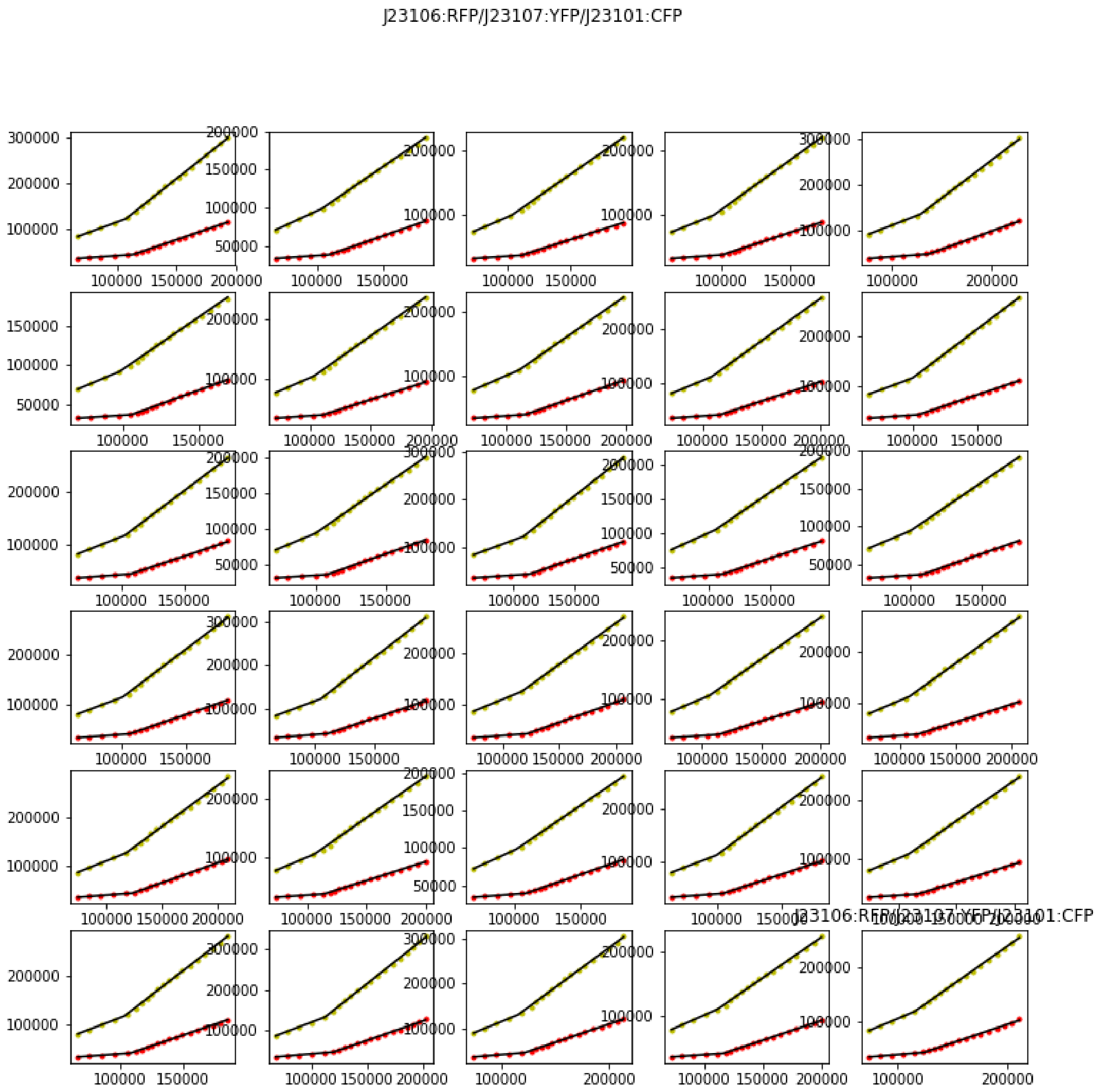


Figure S8. All phase space trajectories for plasmid J23106:RFP/J23107:YFP/J23101:CFP.

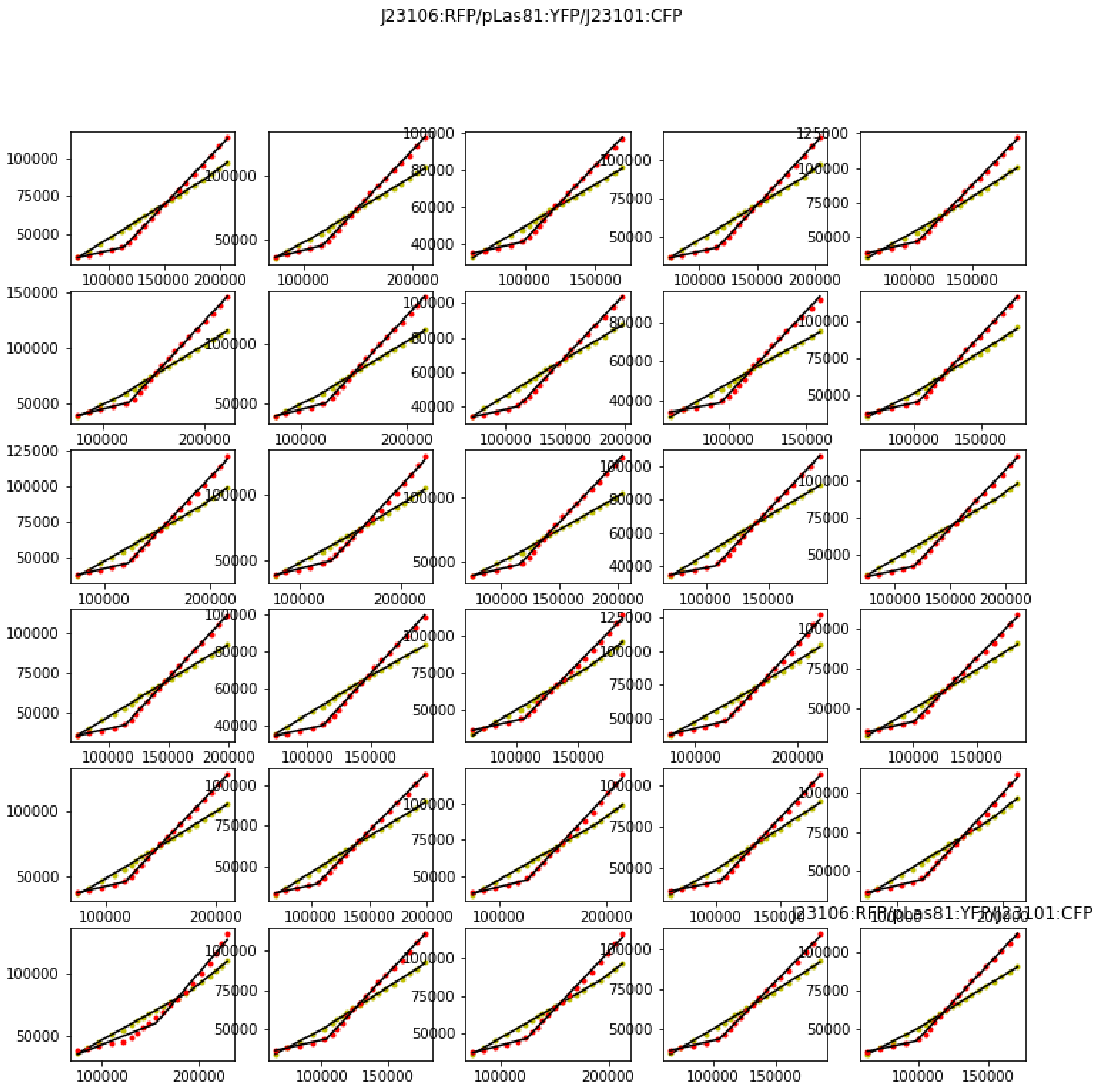


Figure S9. All phase space trajectories for plasmid J23106:RFP/pLas81:YFP/J23101:CFP.

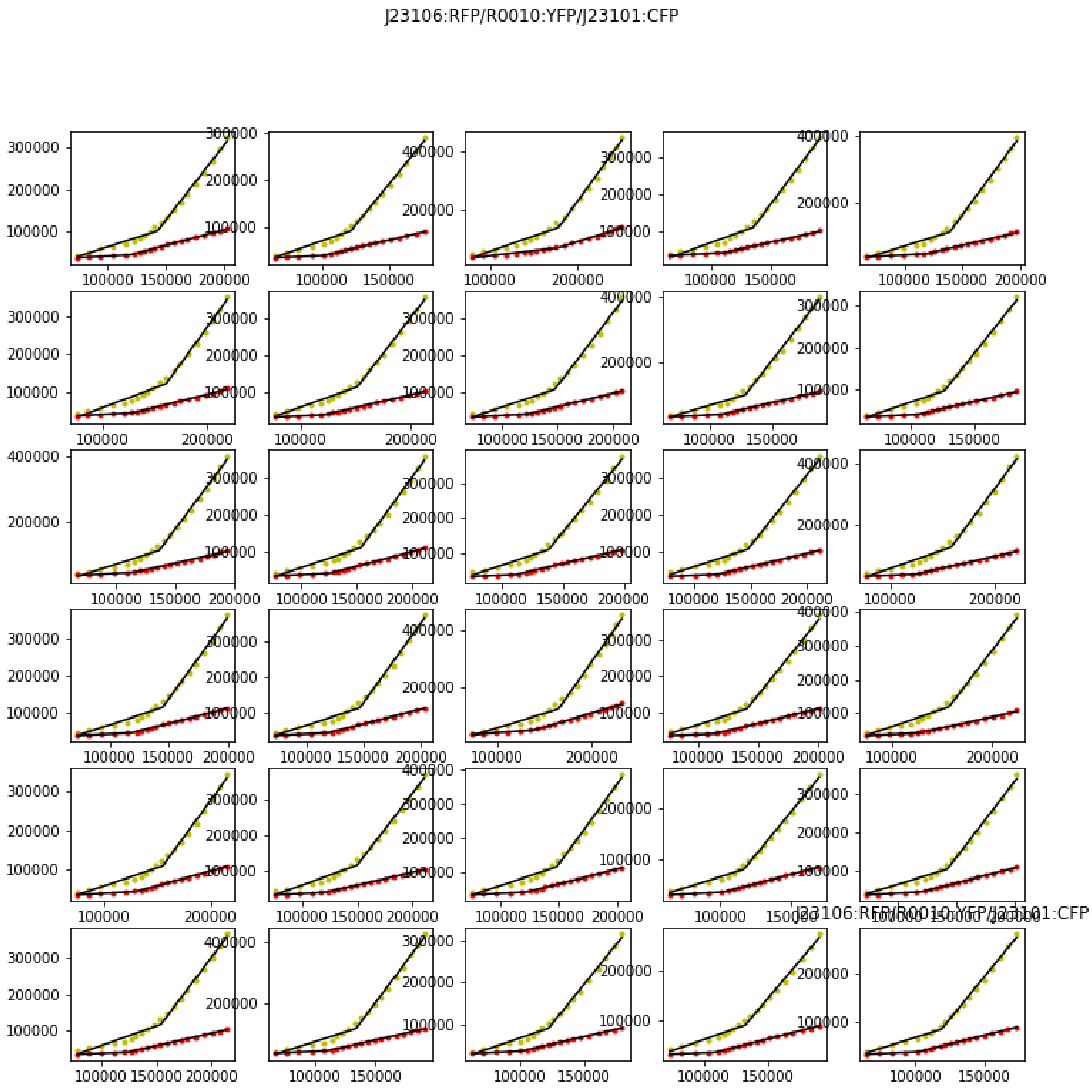


Figure S10. All phase space trajectories for plasmid J23106:RFP/R0010:YFP/J23101:CFP.

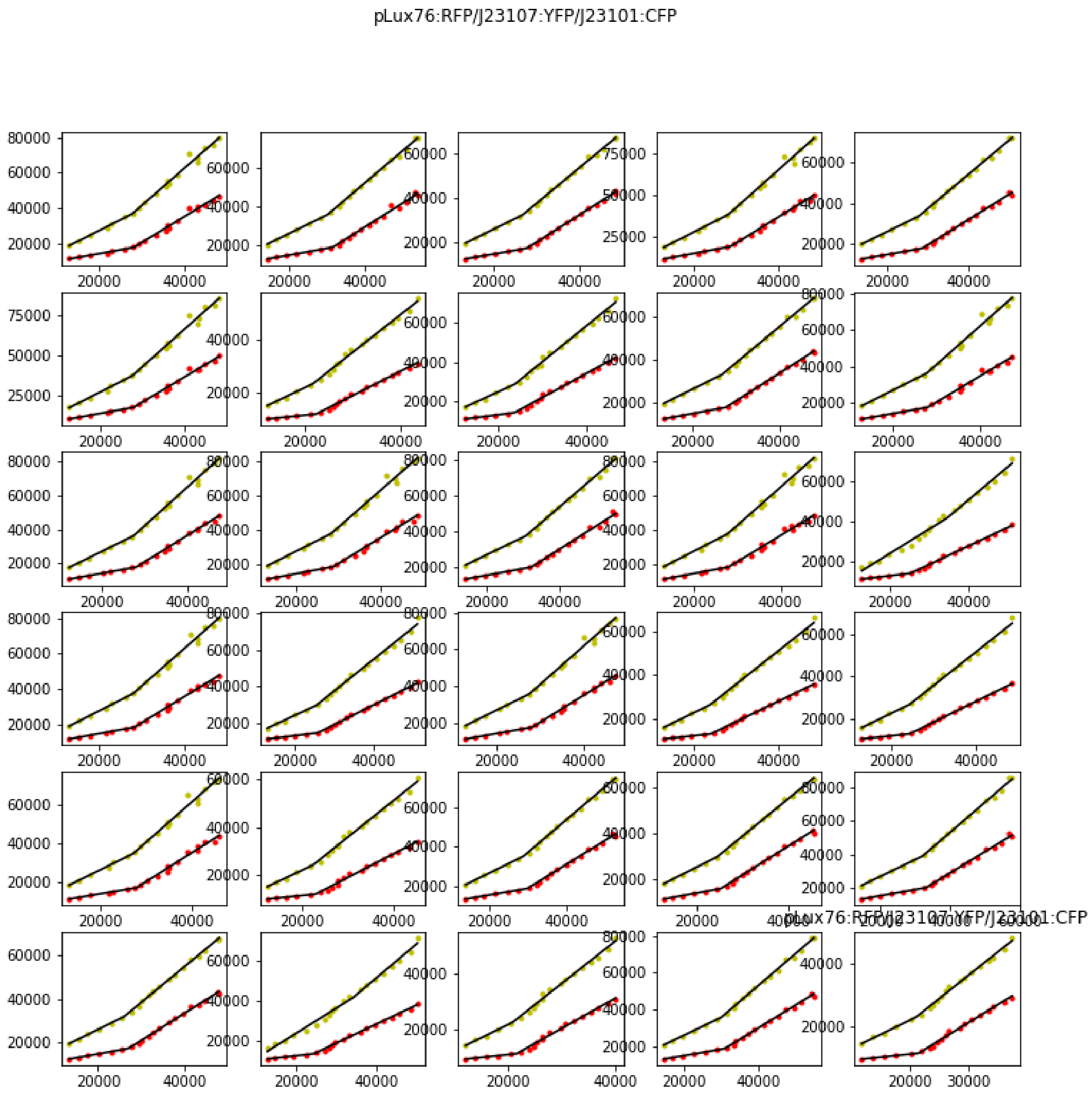


Figure S11. All phase space trajectories for plasmid pLux76:RFP/J23107:YFP/J23101:CFP.

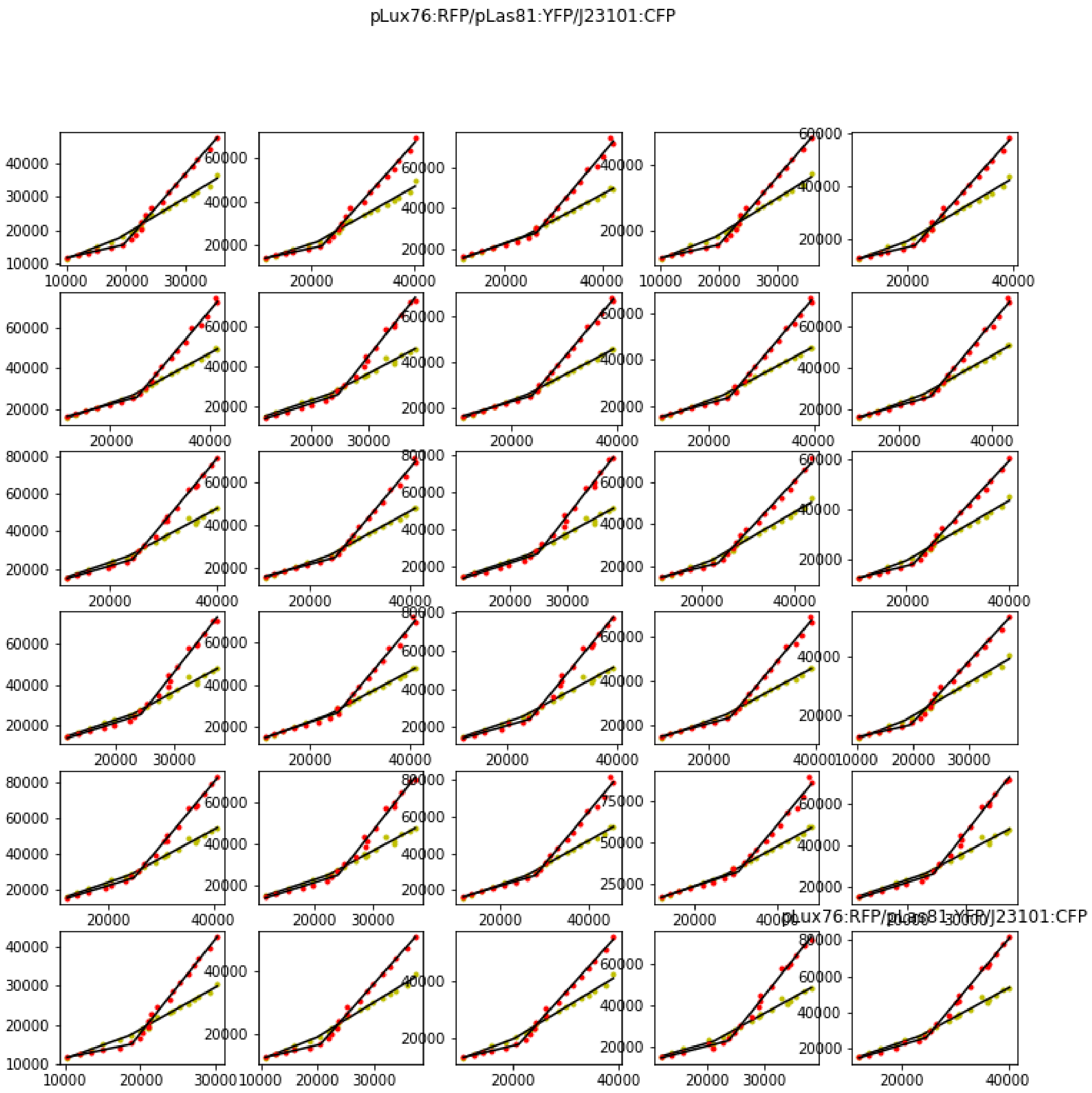


Figure S12. All phase space trajectories for plasmid pLux76:RFP/pLas81:YFP/J23101:CFP.

pLux76:RFP/R0040:YFP/J23101:CFP

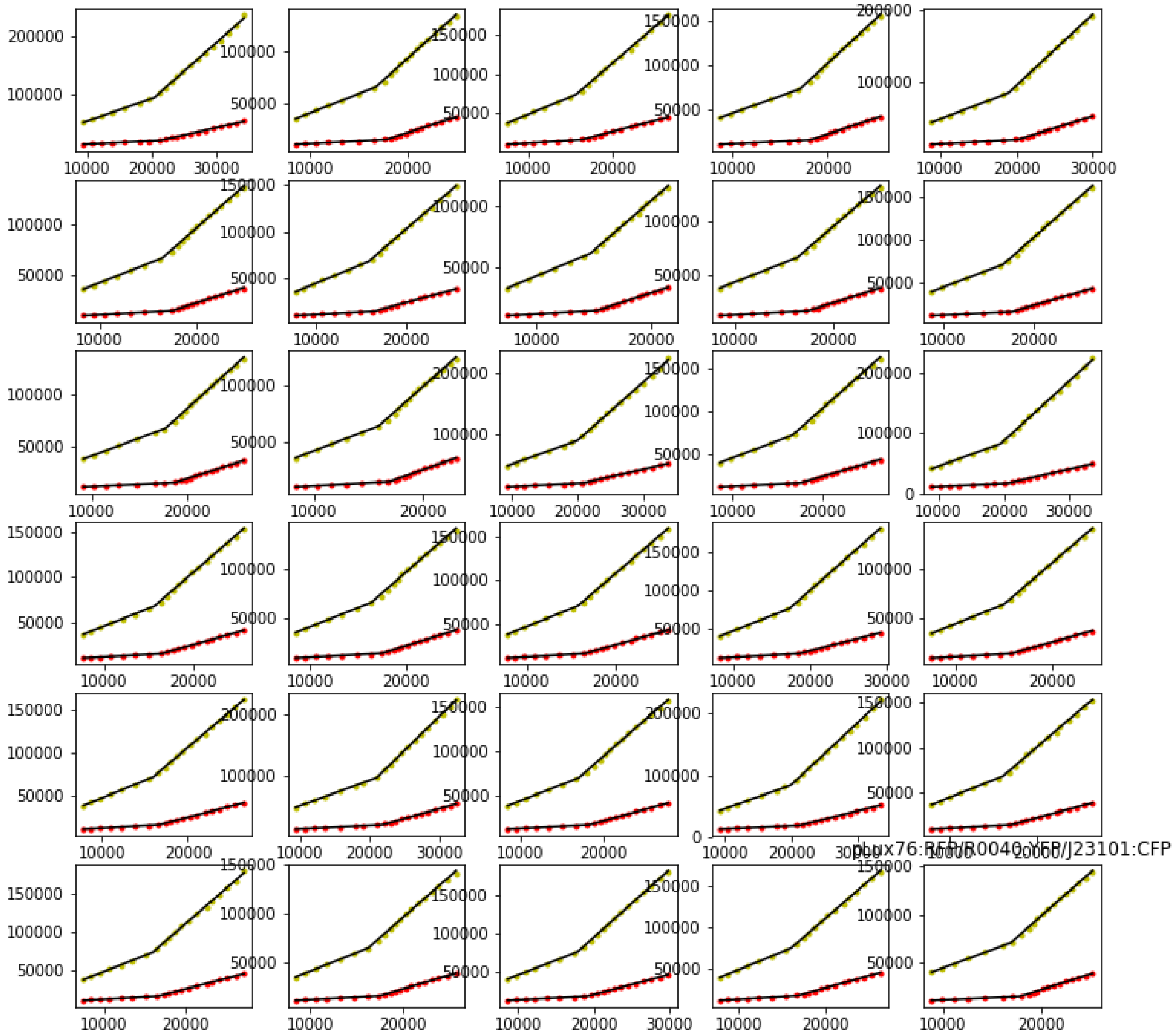

Figure S13. All phase space trajectories for plasmid pLux76:RFP/R0040:YFP/J23101:CFP.

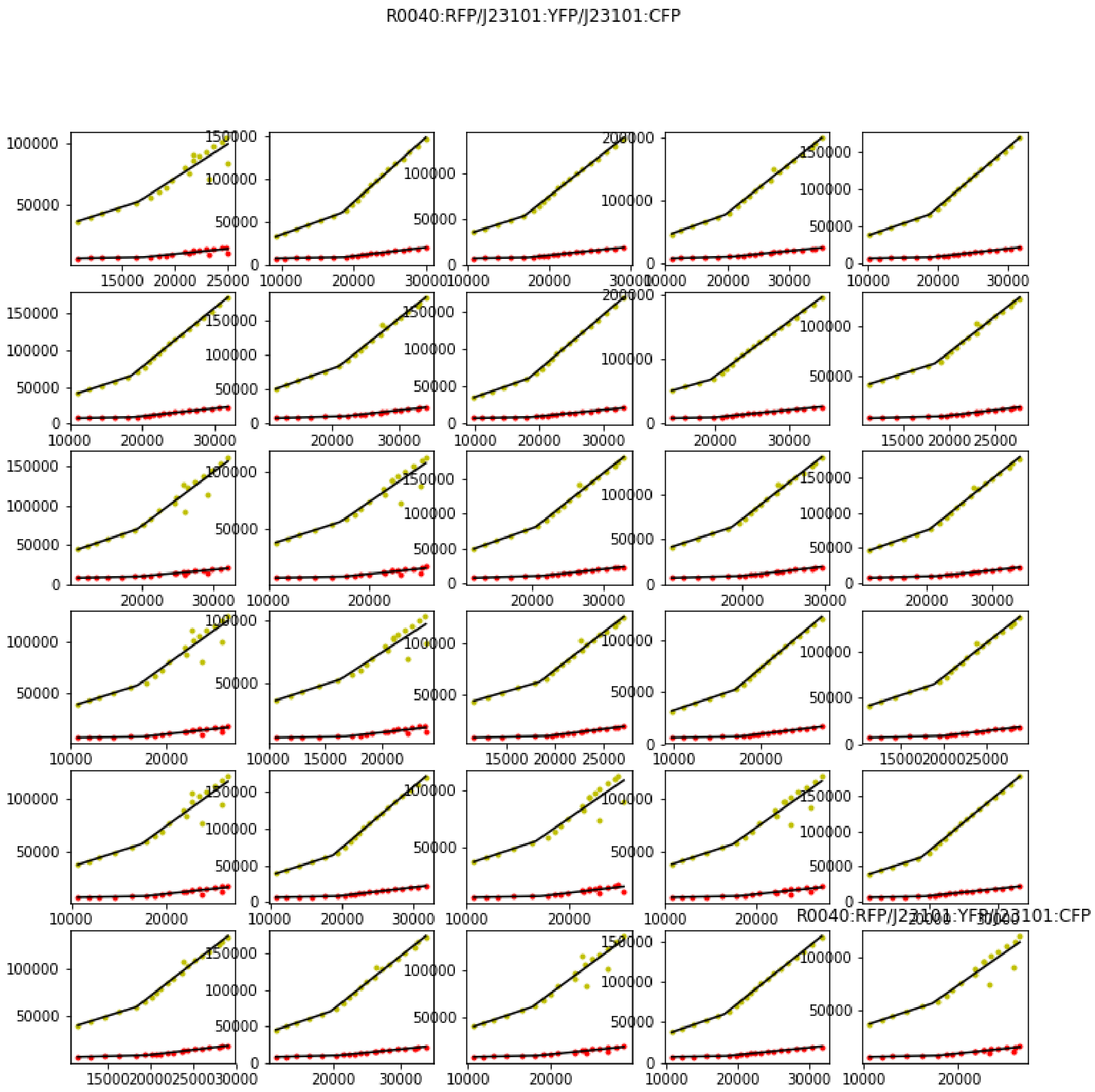


Figure S14. All phase space trajectories for plasmid R0040:RFP/J23101:YFP/J23101:CFP.

R0040:RFP/J23107:YFP/J23101:CFP

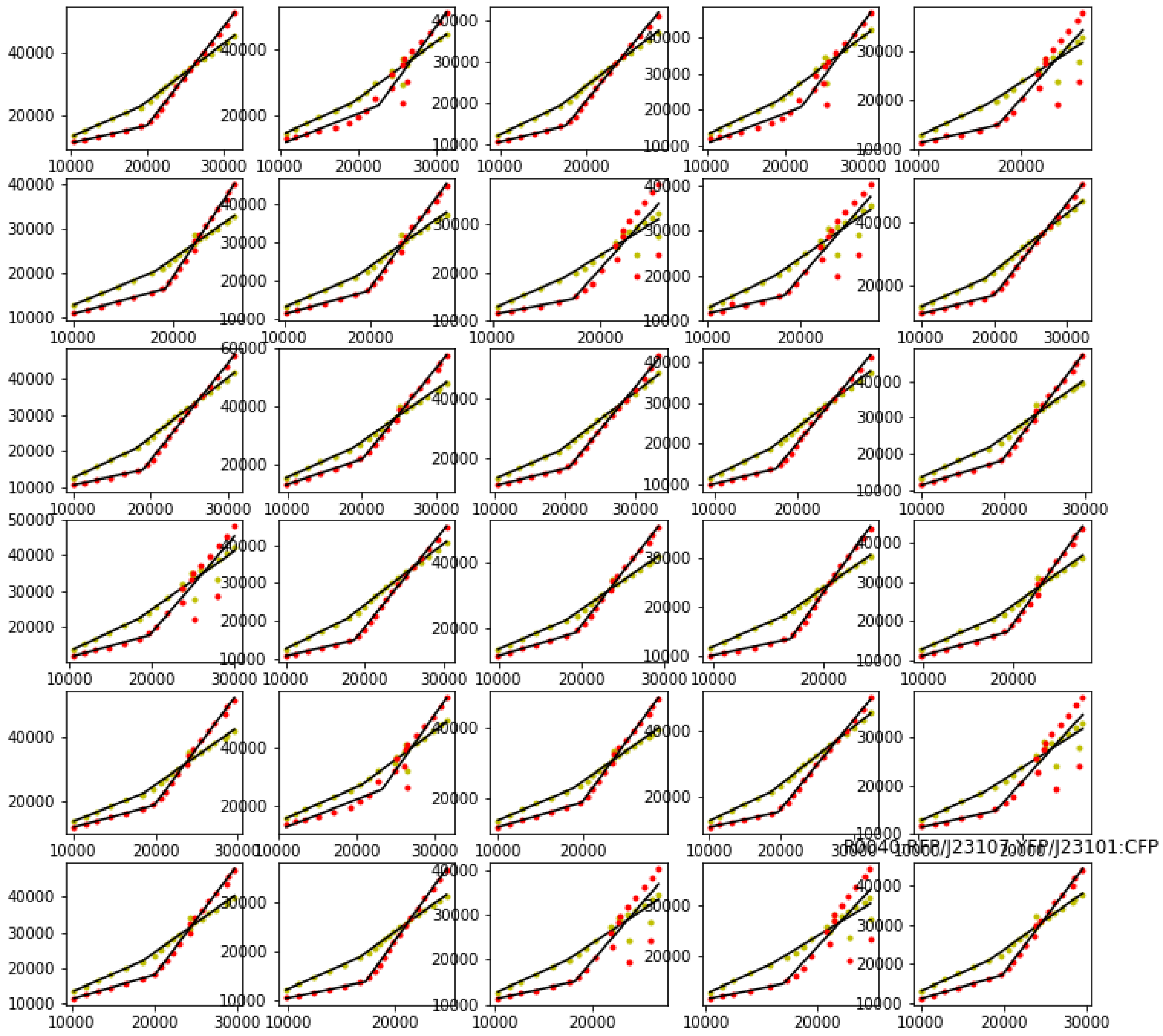

Figure S15. All phase space trajectories for plasmid R0040:RFP/J23107:YFP/J23101:CFP.

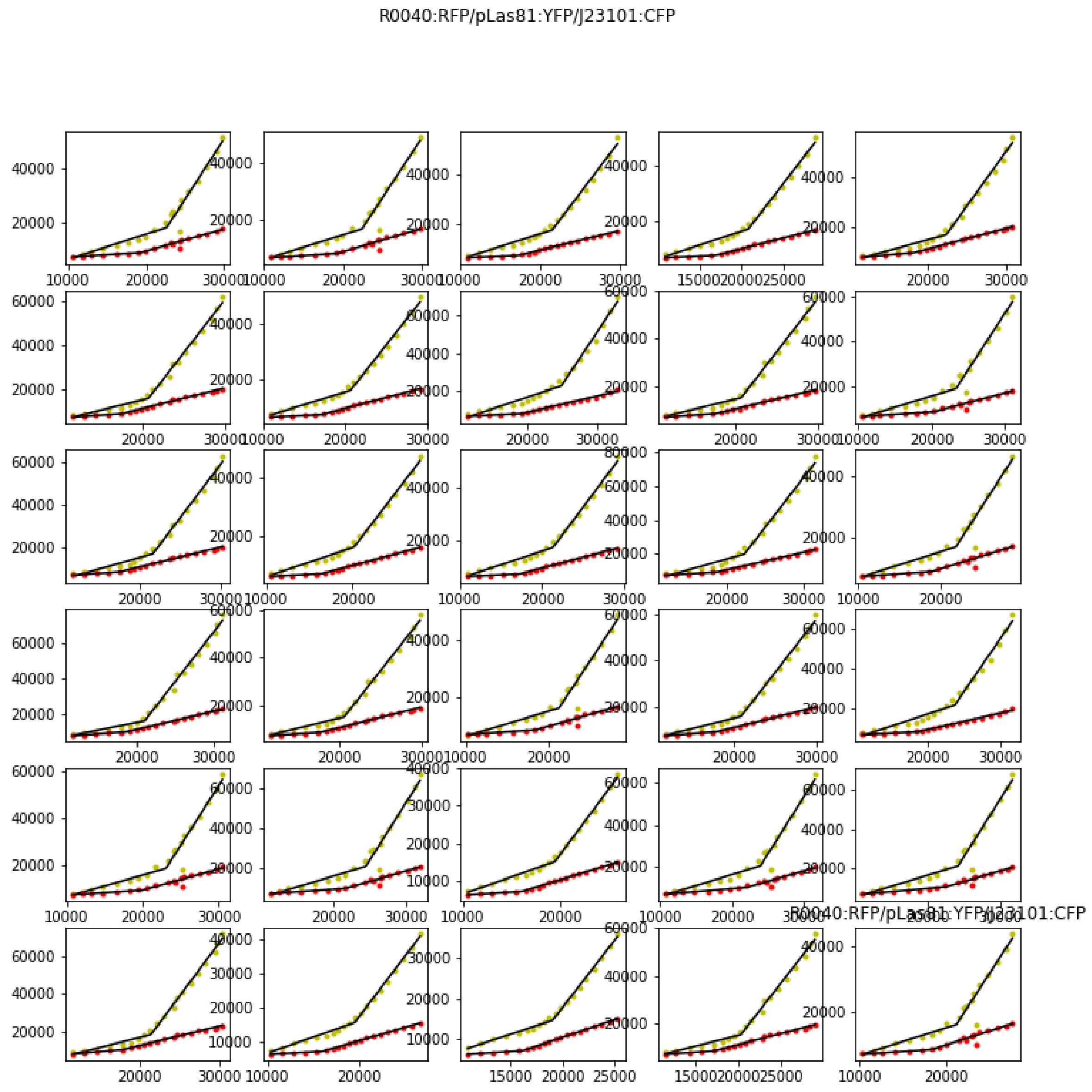

Figure S16. All phase space trajectories for plasmid R0040:RFP/pLas81:YFP/J23101:CFP.

R0040:RFP/R0010:YFP/J23101:CFP

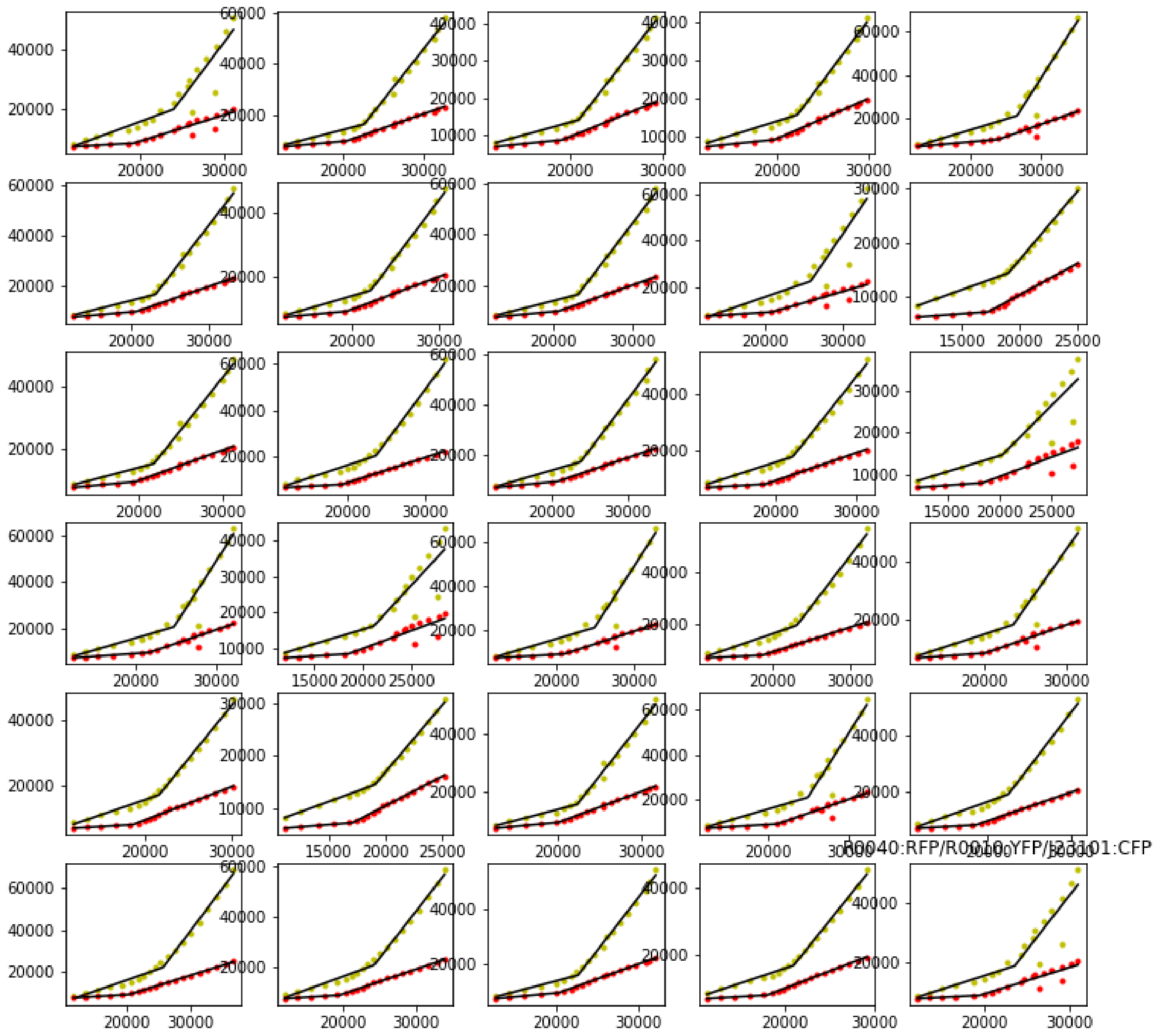

Figure S17. All phase space trajectories for plasmid R0040:RFP/R0010:YFP/J23101:CFP.

**Three TUs Plasmids diagrams**

The following diagrams show the position of the TUs in the assembled plasmids and the parts used in each case. The replication origin and the Kanamycin resistance is shown.

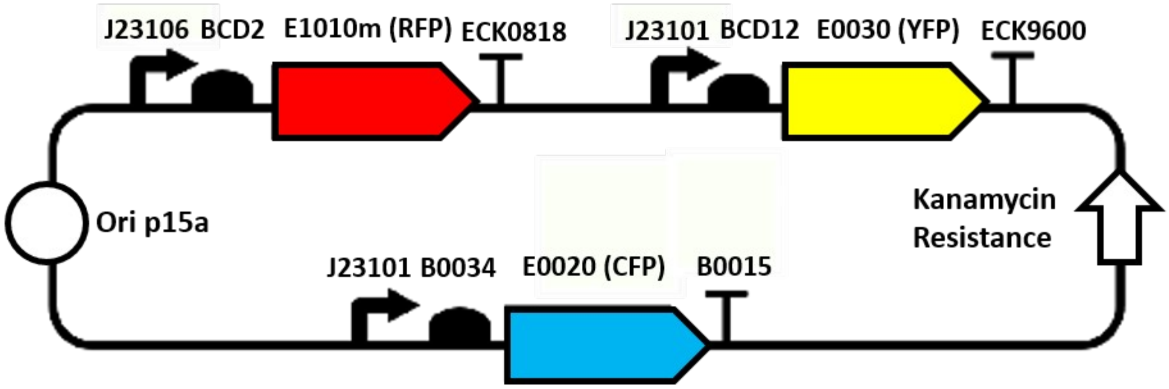


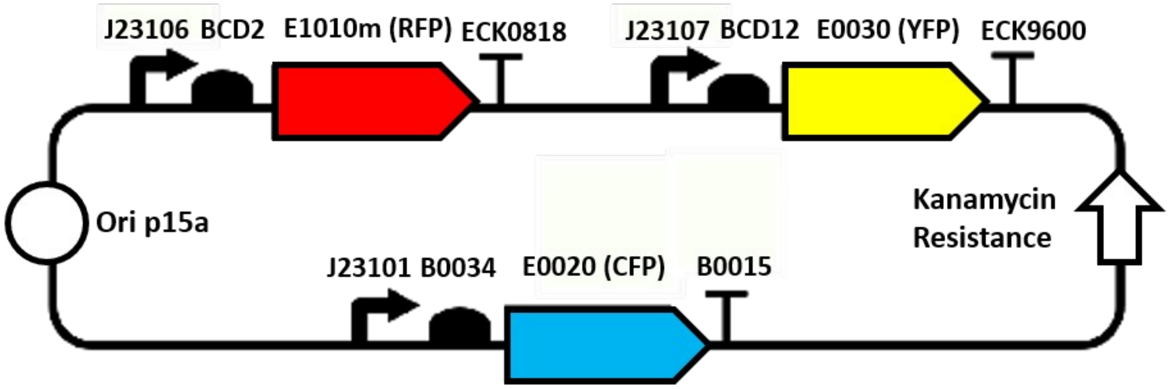

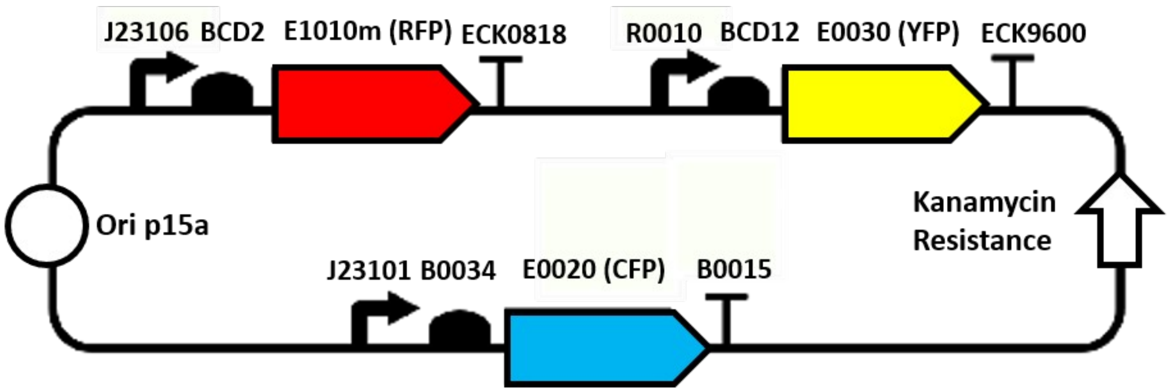
J23106 BCD2 E1010m (RFP) ECK0818
R0010 BCD12 E0030 (YFP) ECK9600
Ori p15a
Kanamycin Resistance
J23101 B0034 E0020 (CFP) B0015

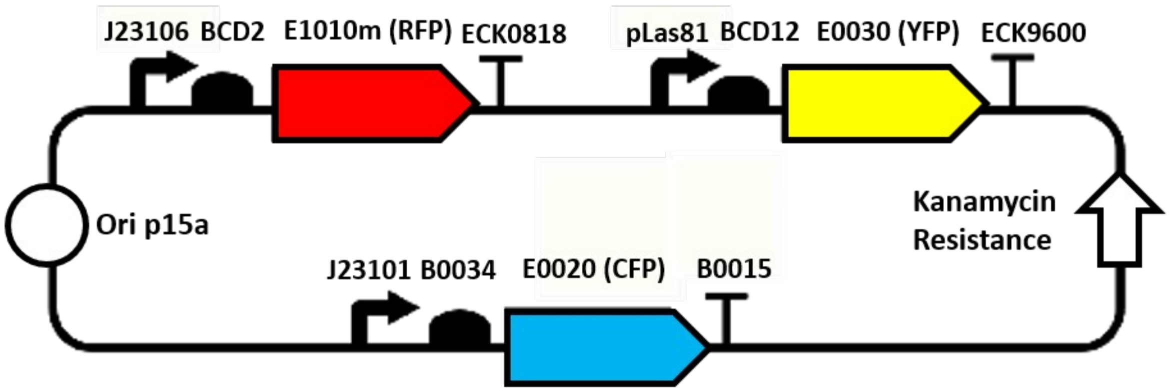
J23106 BCD2 E1010m (RFP) ECK0818
pLas81 BCD12 E0030 (YFP) ECK9600
Ori p15a
Kanamycin Resistance
J23101 B0034 E0020 (CFP) B0015

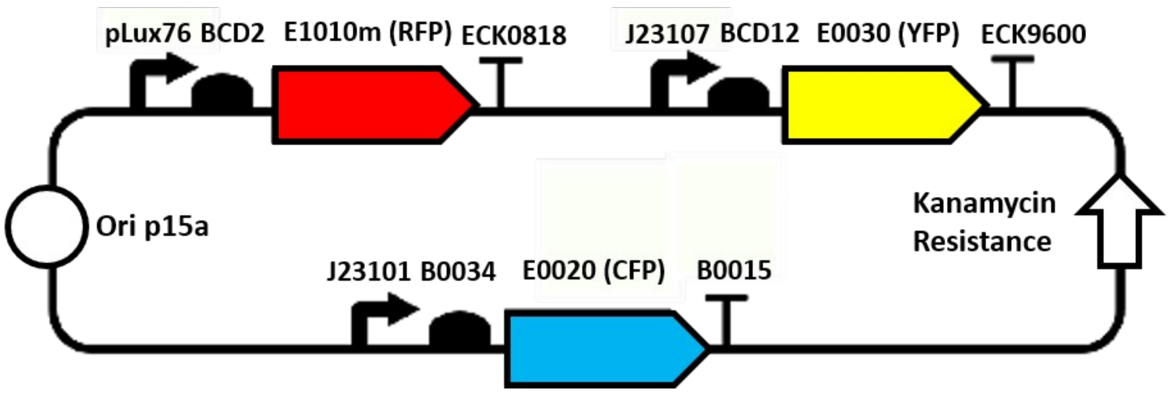
pLux76 BCD2 E1010m (RFP) ECK0818
J23107 BCD12 E0030 (YFP) ECK9600
Ori p15a
Kanamycin Resistance
J23101 B0034 E0020 (CFP) B0015

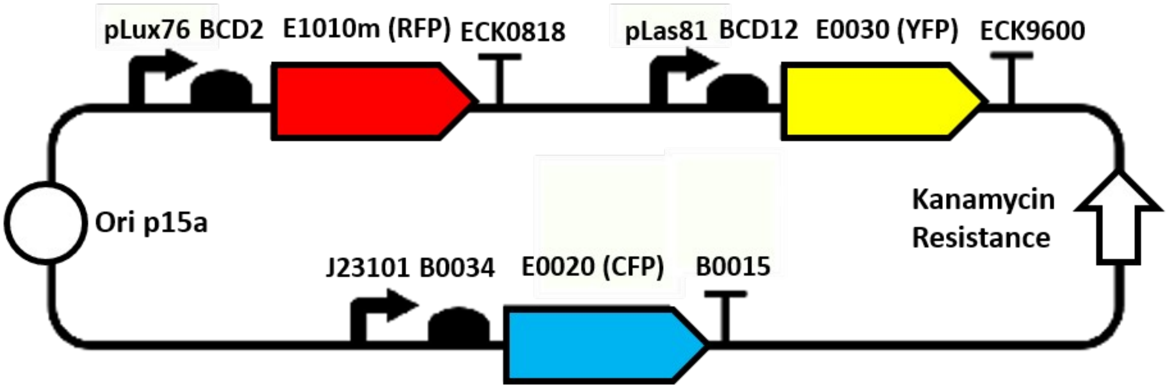
pLux76
BCD2
E1010m (RFP)
ECK0818
pLas81
BCD12
E0030 (YFP)
ECK9600
Ori p15a
Kanamycin Resistance
J23101
B0034
E0020 (CFP)
B0015

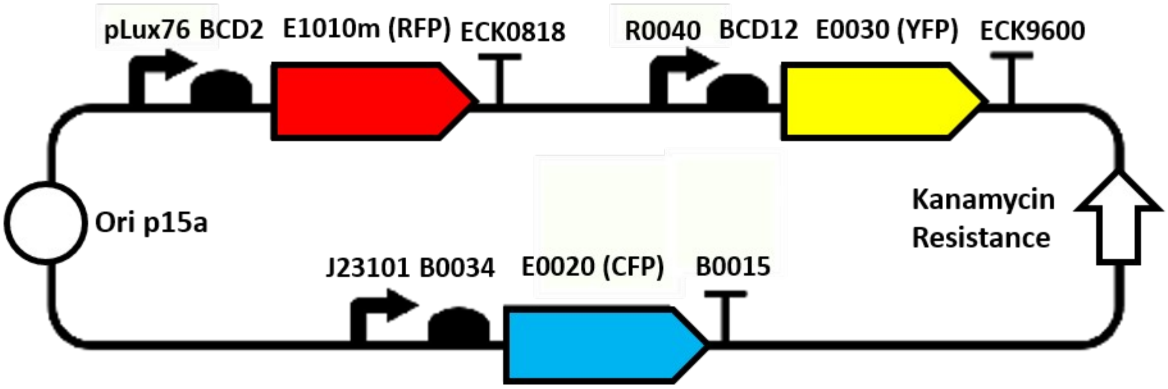
pLux76
BCD2
E1010m (RFP)
ECK0818
R0040
BCD12
E0030 (YFP)
ECK9600
Ori p15a
Kanamycin Resistance
J23101
B0034
E0020 (CFP)
B0015

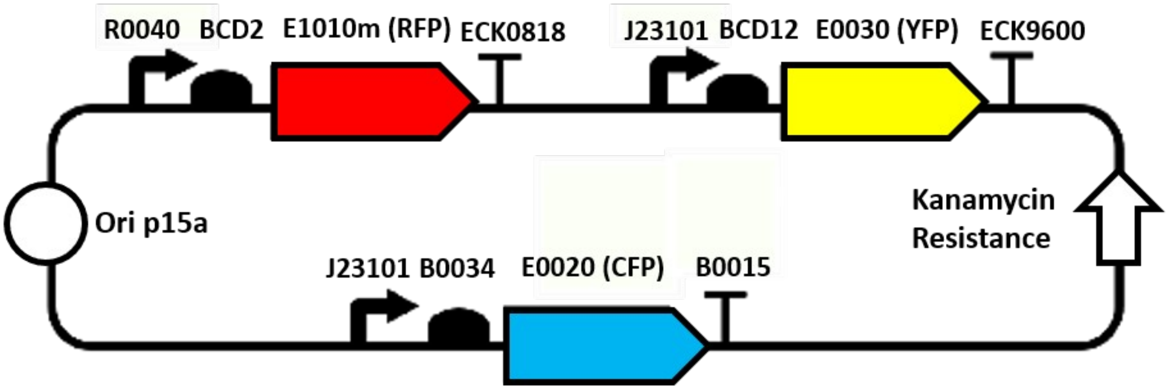
R0040
BCD2
E1010m (RFP)
ECK0818
J23101
BCD12
E0030 (YFP)
ECK9600
Ori p15a
Kanamycin Resistance
J23101
B0034
E0020 (CFP)
B0015

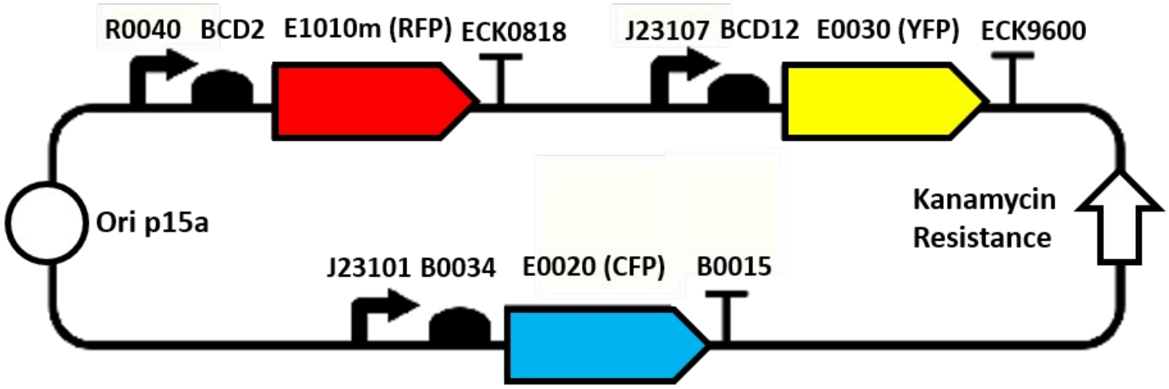
R0040
BCD2
E1010m (RFP)
ECK0818
J23107
BCD12
E0030 (YFP)
ECK9600
Ori p15a
Kanamycin Resistance
J23101
B0034
E0020 (CFP)
B0015

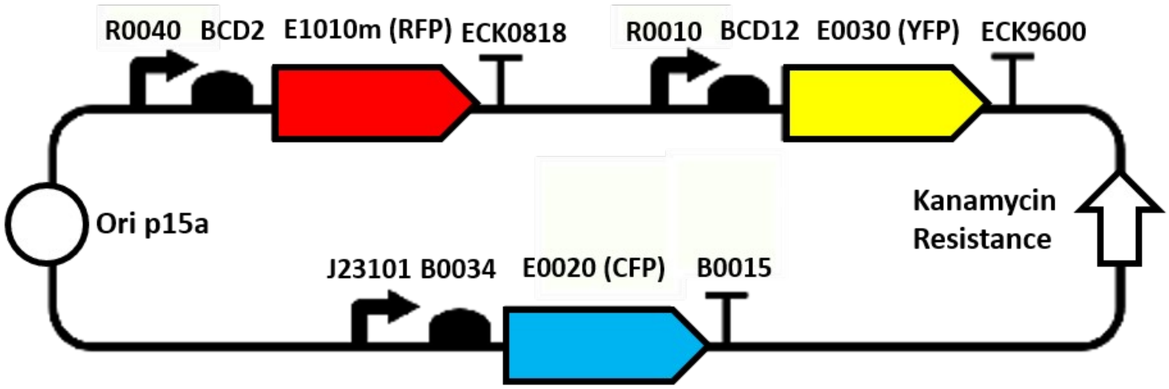
R0040
BCD2
E1010m (RFP)
ECK0818
R0010
BCD12
E0030 (YFP)
ECK9600
Ori p15a
Kanamycin Resistance
J23101
B0034
E0020 (CFP)
B0015

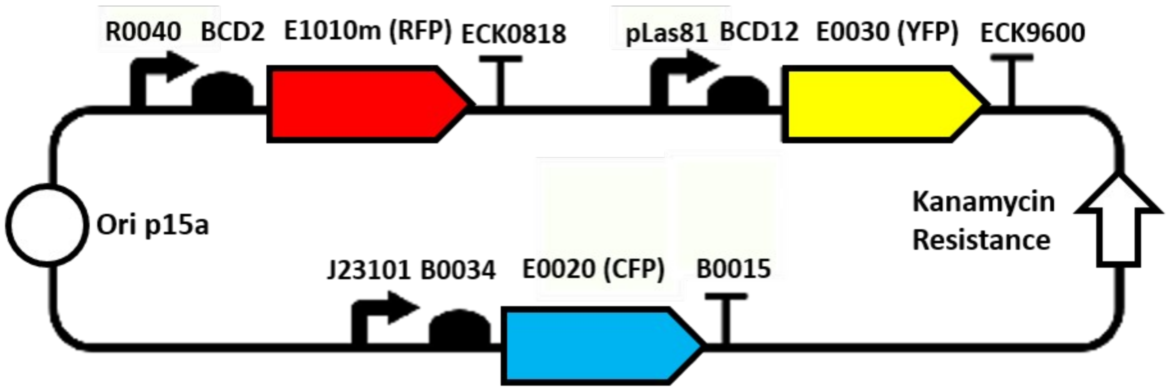
R0040
BCD2
E1010m (RFP)
ECK0818
pLas81
BCD12
E0030 (YFP)
ECK9600
Ori p15a
Kanamycin Resistance
J23101
B0034
E0020 (CFP)
B0015

(9) Oliphant, T. {NumPy}: A guide to {NumPy}.

(10) Jones, E., Oliphant, T., Peterson, P., and others. {SciPy}: Open source scientific tools for {Python}.

(11) Hunter, J. D. (2007) Matplotlib: A 2D Graphics Environment. *Comput. Sci. Eng. 9*, 90–95.

(12) McKinney, W. (2010) Data Structures for Statistical Computing in Python, in *Proceedings of the 9th Python in Science Conference* (van der Walt, S., and Millman, J., Eds.), pp 51–56.

(13) Kruskal, W. H., and Wallis, W. A. (1952) Use of ranks in one-criterion variance analysis. *J. Am. Stat. Assoc. 47*, 583–621.
